# Combining ^13^C, ^15^N, and ^2^H to measure feeding and metabolic activity in marine, shal-low-water sponges – A pilot study

**DOI:** 10.1101/2024.11.27.625585

**Authors:** Tanja Stratmann, Diana X. Sahonero-Canavesi, Marcel T. J. van der Meer

## Abstract

**Background:** Shallow-water sponges can reach high densities and have several functions in the ecosystem, such as, providing microhabitats for other species or being involved in benthic-pelagic coupling. They also have a fast cell turnover so they can serve as test animals for the development of new methods to measure the metabolic activity of individual organisms. Here, we measured the feeding and metabolic activity of the common intertidal sponge *Halichondria panicea* using a triple stable isotope labeling experiment with a ^13^C and ^15^N-enriched bacteria as substrate and ^2^H from deuterated water. We also constrained pathways of phospholipid-derived fatty acid (PLFA) biosynthesis in the sponge and its microbiome.

**Methods:** Sponges were collected in the Eastern Scheldt (North Sea) and incubated in 1% ^2^H-enriched seawater in the presence of ^13^C- and ^15^N-enriched substrate of inactivated bacteria for 12 h. Water samples for the analysis of dissolved inorganic carbon (DIC), ^13^C-DIC, and inorganic nutrients were taken at the begin and the end of the incubations, while oxygen concentration in the water was recorded continuously. Seawater incubations with the addition of substrate bacteria served as blanks and dead sponges incubated in 1% ^2^H-enriched seawater served as controls for the incorporation of ^2^H into inactive tissue. At the end of the experiment, sponges were sampled for bulk analyses of ^13^C, ^15^N, and ^2^H in sponge tissue, and for ^13^C and ^2^H incorporation into phospholipid-derived fatty acids (PLFAs).

**Results:** Sponges consumed oxygen, nitrite, nitrate, and silicon, while they excreted ammonia and released ^13^C-DIC. They incorporated 4.82 μmol ^13^C mmol C^−1^ d^−1^, 2.46 μmol ^15^N mmol C^−1^ d^−1^, and 0.49 μmol ^2^H mmol C^−1^ d^−1^ into bulk sponge tissue. The PLFAs extracted from the sponges contained on average 5.61 μg ^13^C g^−1^ dry mass (DM) sponge and 5.43 ng ^2^H g^−1^ DM sponge. Most ^13^C was incorporated into bacteria-specific PLFAs derived from the substrate bacteria; ^2^H, in comparison, was mostly built in sponge-specific PLFAs.

**Conclusion:** *H. panicea* from the Eastern Scheldt were in a similar condition as starving specimens from the Baltic Sea which suggests that the large bivalve stocks in the Eastern Scheldt outcompete this sponge for food. During the incubation experiment, however, the sponges were well fed as indicated by the silicon and oxygen uptake rates. The significantly higher uptake rates of ^2^H by living sponges compared to dead sponges proved that deuterated water can be used to measure the metabolic activity of individual filter-feeding specimens, but more test with species of diverse biological traits and growth stages are recommended. Additionally, we suggest to focus on compound-specific ^2^H uptake, as these data were less affected by ^1^H exchange of non-covalently bound ^1^H. Assessing ^13^C- and ^2^H-enriched PLFAs of the incubated *H. panicea* also revealed that the sponge microbiome produced the bacteria-specific PLFAs *i*-C14:0, *ai*-C15:0, *i*-C17:0, and *ai*-C17:0 using C from the substrate bacteria. The *ai*-C15:0 was subsequently elongated and desaturated to build the sponge-specific PLFA *ai*-C25:2. Other sponge-specific PLFAs were formed using C14:0 and C16:0 from the sub-strate bacteria as precursors.

## Introduction

Sponges, phylum Porifera, are prominent members of the benthic community in shallow-water ecosystems. For example, at rocky intertidal sites of the temperate Bay of Brest (NW France, NE Atlantic) sponges had a density of 3.4 ± 2.5 ind. m^−2^, a biovolume of 0.124 ± 0.132 ml m^−2^, and a species richness of siliceous sponges of 4 (López-Acosta et al., 2022). In temperate hard-substrate subtidal areas, sponge densities can reach 16.4 + 9.3 ind. m^−2^ (rocky subtidal, Bay of Brest; (López-Acosta et al., 2022)) to 50 ind. m^−2^ at vertical surfaces and 100 ind. m^−2^ at inclined surfaces (Lough Hyne, Ireland, NE Atlantic; (Bell & Barnes, 2000)). In sandy/ muddy subtidal areas, in comparison, sponge densities at the temperate Bay of Brest were 0.5 ± 1.0 ind. m^−2^ (López-Acosta et al., 2022) and the four dominant sponge species in the tropical Mazatlan Bay (Mexico, E Pacific) had densities of <1.2 ind. m^−2^ (*Mycale (Zygomycale) ramulosa, Callyspongia (Callyspongia) californica, Clathria (Microciona)* sp.) to 1.22 ind. m^−2^ (*Haliclona (Soestella) caerulea*) (Ávila et al., 2011).

Sponges also have important functions in the temperate marine shallow-water ecosystem, like serving as “agents of biological disturbance” (Bell, 2008) by competing for space (Khalaman & Komendantov, 2016), enhancing benthic-pelagic coupling (Perea-Blázquez, Davy & Bell, 2012; Perea-Blázquez et al., 2013) and cycling of nutrients (López-Acosta et al., 2022), serving as prey for spongivores (Ruzicka & Gleason, 2009), and providing microhabitats and increasing local biodiversity (Abdo, 2007; Chin et al., 2020). In fact, in SW Australia (Indian Ocean), two temperate sponge species *Haliclona* sp. 1 (green morphotype) and *Haliclona* sp. 2 (brown morphotype) hosted 24 taxa of endofauna belonging to 5 different phyla which had a total endofauna biomass of 101 g wet mass (WM; *Haliclona* sp. 1) and 55 g WM (*Haliclona* sp. 2) (Abdo, 2007).

The intertidal sponge *Halichondria (Halichondria) panicea* (Pallas, 1766), commonly known as the breadcrumb sponge, has a cosmopolitan distribution ranging from the Arctic (e.g. Svalbard) to the tropics (e.g. Sri Lanka) with a frequent occurrence in the NE Atlantic and the NE Pacific (OBIS, 2024). *H. panicea* grows as epibiont on top of red algae (Barthel, 1986), blue mussels (Khalaman & Komendantov, 2011), ascidians (Khalaman & Komendantov, 2011), scallops (Forester, 1979), and oysters (Stratmann, personal observation), or it uses boulders as hard substrate (Thomassen & Riisgård, 1995). This sponge has a seasonal growth cycle which is partly linked to changes in water temperature, with low growth in spring, maximum growth in summer, and a decline in late summer to early autumn, (Barthel, 1986), when it releases its larvae (Amano, 1986). *H. panicea* is a suspension feeder that filters phytoplankton (Riisgård et al., 1993; Riisgård, Kumala & Charitonidou, 2016; Lüskow et al., 2019) and bacteria (Lüskow et al., 2019) out of the water column with filtration rates of 31.0 to 46.1 ml water g^−1^ dry mass (DM) sponge min^−1^ (Riisgård et al., 1993; Riisgård, Kumala & Charitonidou, 2016).

Sponges can have a particularly fast cell turnover (i.e., the balance between proliferated and lost cells (Alexander et al., 2014)): The encrusting tropical sponge *Halisarca caerulea* Vacelet & Donadey, 1987, for instance, seems to use a large amount of the carbon (C) it consumes to renew the choanocytes in its tissue (de Goeij et al., 2009) with a proliferation rate of choanocytes of 17.6 ± 3.3% in 6 h (Alexander et al., 2014). The mangrove sponge *Mycale* (*Carmia*) *microsigmatosa* Arndt, 1927 has even a choanocyte proliferation rate of 70.5 ± 6.6% and a mesohyl cell proliferation rate of 19.5 ± 4.9% in 6 h (Alexander et al., 2014). These proliferation rates were measured by labelling with 5-bromo-2′-deoxyuridine (BrdU) which is incorporated into the DNA during the S-phase of the cell cycle (Nowakowski, Lewin & Miller, 1989). Another approach to measure cell proliferation rates is labelling with deuterated water (^2^H_2_O, known as deuterium oxide or heavy water) (Foletta et al., 2016).

Deuterated water has been used to trace the uptake of water in terrestrial plants (e.g. (Kowaljow & Fernández, 2011; Kulmatiski & Beard, 2013; Huo et al., 2020; Aguzzoni et al., 2022; Giuliani et al., 2023)), to study the biosynthesis of lipids (e.g. (Kellermann et al., 2016; Zhang et al., 2019; Pilecky et al., 2022)), to assess the energy metabolism and expenditure of invertebrates and vertebrates (e.g. (Qiu, Lacey & Bedding, 2000; Junghans et al., 2018)), and to track the temporal and spatial distribution of mosquitos (Faiman et al., 2019). Here, we use deuterated water to determine the metabolic activity of sponges, whereupon we define metabolic activity as “the chemical changes that occur in a living animal […] as a result of metabolism” (Park, 2012) including ‘synthetic reactions’ like the production of proteins and fatty acids and ‘destructive reactions’ like the split of sugars into CO_2_ and water (Park, 2012).

Fatty acids are part of complex polar lipids and building blocks of membranes (van Deenen, 1966; Spector & Yorek, 1985). They play a role as energy source (Lindsay, 1975), are involved in gene expression regulation (Sampath & Ntambi, 2004), and in signal transduction (Graber, Sumida & Nunez, 1994; Faergeman & Knudsen, 1997). Several fatty acids can only be produced by specific groups of organisms. For instance, iso (*i*) and ante-iso (*ai*)-branched C chain fatty acids are produced by bacteria (e.g., *i*-C15:0, *ai*-C15:0, *i*-C17:0, *ai*-C17:0 (Kaneda, 1991)), whereas sponge-specific long chain fatty acids (≥C24) are produced by sponges via elongation of shorter chain fatty acids (Carballeira et al., 1986; de Kluijver et al., 2021). Therefore, measuring the incorporation of ^2^H into bacteria- and sponge-specific fatty acids should provide information about the metabolic activity of the sponge and its associated microbial community (i.e., the sponge microbiome).

The aim of this study was to measure the feeding activity of *H. panicea* and to better understand the potential biosynthetic pathways of fatty acid production involved. We further wanted to test whether ^2^H_2_O could be used as substrate-independent tracer to study metabolic activity of filter feeders.

## Materials and Methods

### Study site and sampling location

The Eastern Scheldt (Oosterschelde) is a semi-enclosed 350 km^2^ large tidal bay in the south-western part of the North Sea with a tidal range of 2.9 to 3.5 m (Jiang et al., 2019) and a tidal exchange of ∼20,000 m^3^ s^−1^ (Jiang, Soetaert & Gerkema, 2019). In the central Eastern Scheldt the turnover time of seawater is 90 to 120 days (Jiang, Soetaert & Gerkema, 2019) and the residence time is about 100 days (Jiang et al., 2019). This bay harbors extensive shellfish cultures and fisheries with a focus on cockles (*Cerastoderma edule* (Linnaeus, 1758)), mussels (*Mytilus edulis* Linnaeus, 1758), and non-native Pacific oysters (*Magallana gigas* (Thunberg, 1793)) (Smaal et al., 1986, 2013; Smaal & Lucas, 2000; Smaal, Kater & Wijsman, 2009).

Specimens of the sponge *H. panicea* (Fig. 1) were collected during low tide at the Sint Pieters Polder intertidal flat (51.4729°N, 4.0928°E) in the central Eastern Scheldt in mid-August 2019. The tidal range at this intertidal flat is 1.5 m and it is exposed to air for approximately 6.5 h per tidal cycle (12.4 h).

**Figure 1.**
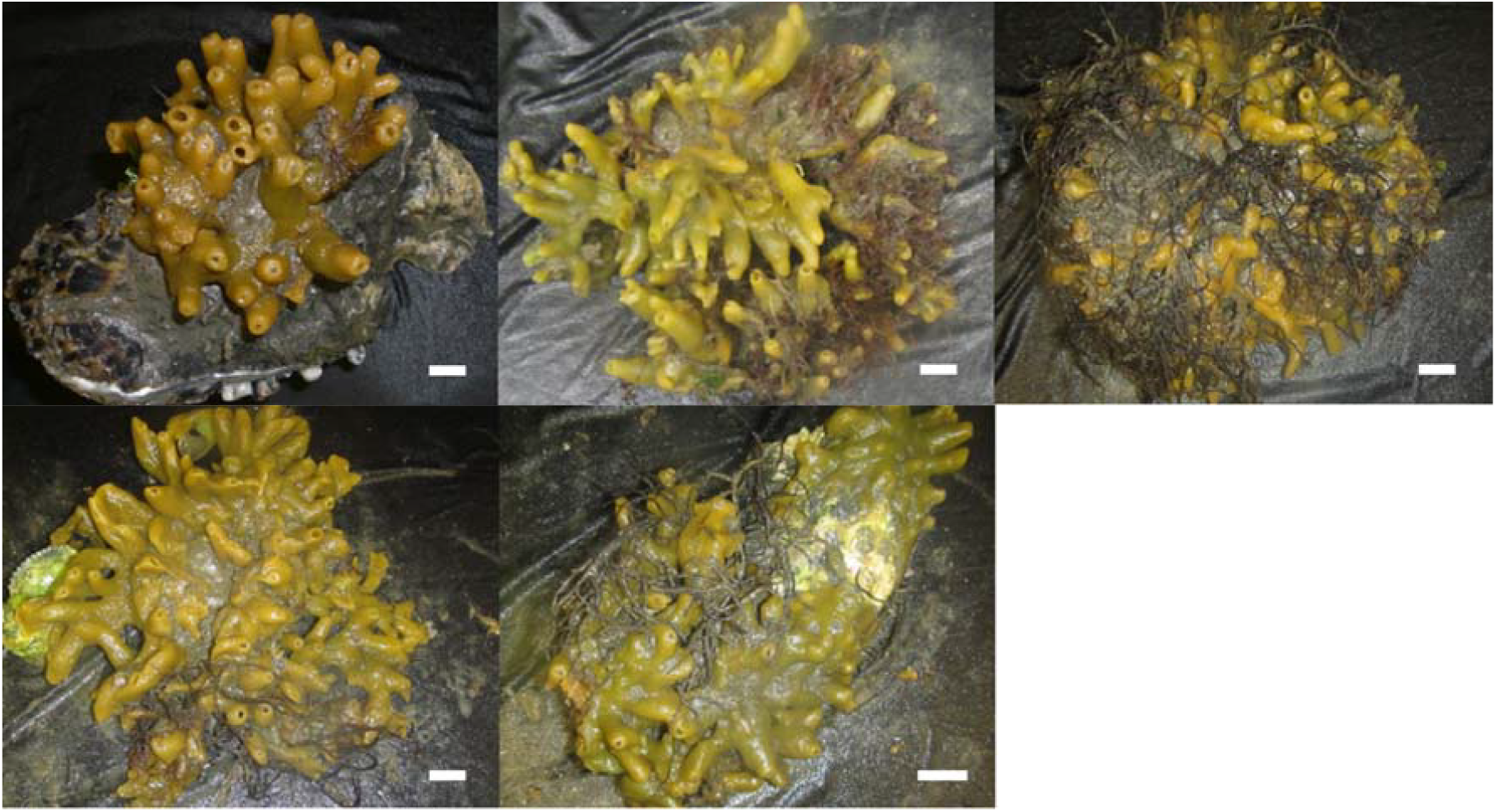
All sponge specimens that were incubated in this experiment. White bar represents 1 cm length.

### Preparation of labelled substrate bacteria

Labelled substrate bacteria was prepared as described in Mueller et al., (2014): A few ml seawater from the Eastern Scheldt with its natural, pelagic bacteria community were added to 1 l culture medium. The culture medium was a variation of the M63 minimum medium (Miller, 1972; Table S1) whereupon (NH_4_)_2_SO_4_ was replaced by Na_2_SO_4_ and NH_4_Cl. 50 atom% of this NH_4_Cl consisted of ^15^NH_4_Cl (99% atom-^15^N; Euroisotop, France) and 30 atom% of the glucose consisted of ^13^C D-glucose (99% atom-^13^C, Cambridge Isotope Laboratories, USA), which was added in four time steps. After 4 d dark incubation at room temperature, the bacteria solution (cell concentration: 80,000 cells ml^−1^) was concentrated by centrifugation, rinsed with artificial seawater to remove any residuals of the labelled substrate, and kept frozen (−20°C) until the start of the experiment. The bacteria concentrate contained 2 10^11^ cells ml^−1^ and had an organic (org.) C content of (mean ± std) 22.1 ± 1.11% (28.6±0.07 at% ^13^C; n = 5) and a total nitrogen (TN) content of 4.66 ± 0.23% (51.0 ± 0.31 at% ^15^N).

### Experimental set-up and procedure

This study consisted of the main treatment (i.e., incubation of alive sponges) and two control treatments (i.e., seawater incubation, incubation of dead sponges) which all were subject to a series of manipulations and measurements (Fig. 2).

**Figure 2.**
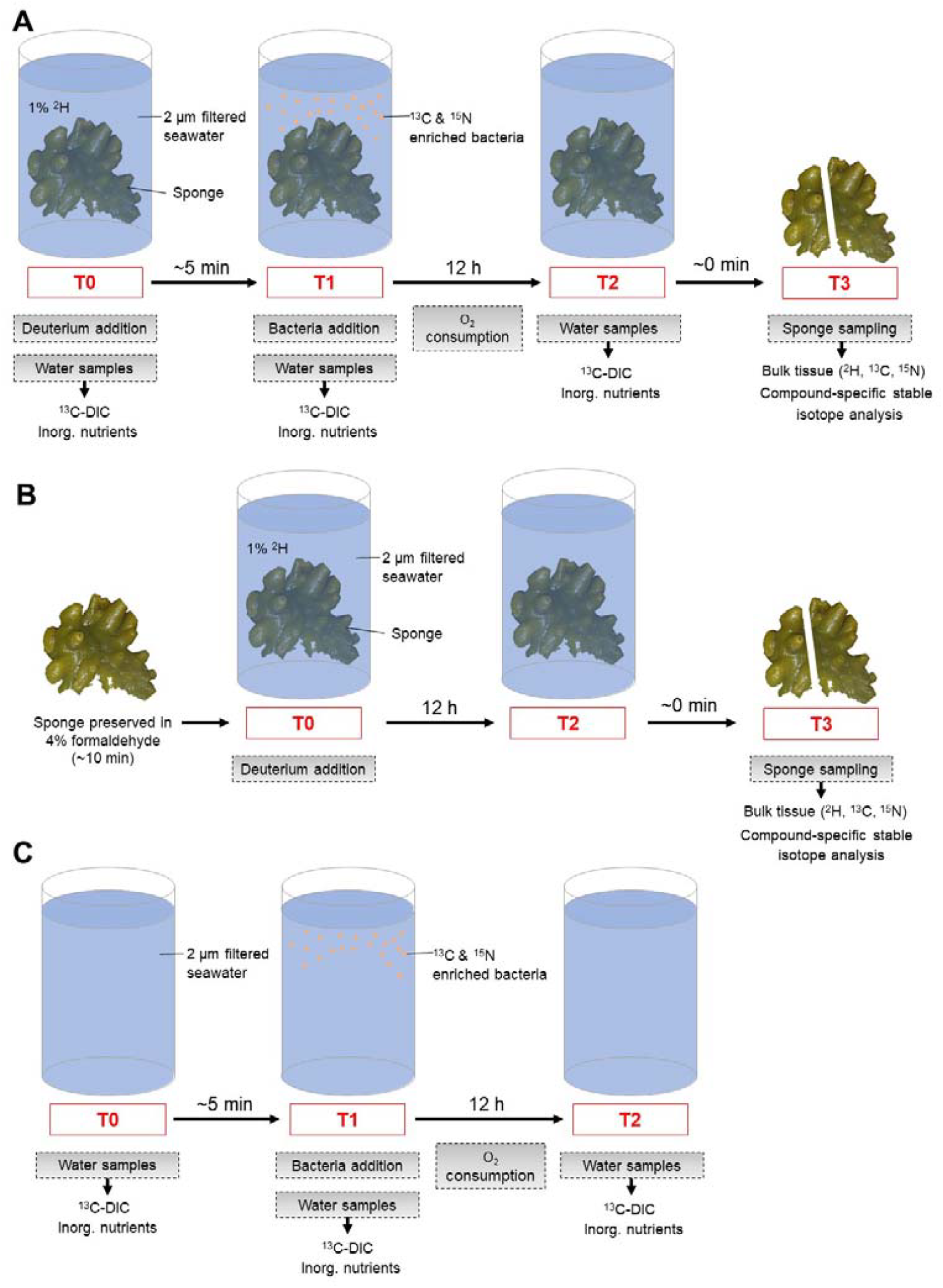
Scheme of individual manipulation and sampling steps during the (A) alive sponge incubation, (B) dead sponge incubations, and (C) the seawater incubation. “T” indicates each sampling step and grey boxes describe the manipulations and measurements conducted at each step. The arrow specifies which variable was analyzed.

Within 2 h after collection, sponges for the main treatment (Fig. 2A) were transported to a climate room in the facilities of NIOZ-Yerseke (The Netherlands). There, sponges were gently washed with pre-filtered (2 μm pore size) seawater from the Eastern Scheldt to remove attached sediment. Five cleaned sponges were assigned individually to incubation chambers (V = 8.5 l) that were filled to 1/3 with pre-filtered seawater. After adding 85 ml ^2^H-water (99.9% ^2^H_2_O, Eurisotop, France; equivalent to 1% ^2^H-enrichment), all chambers were filled to the top with pre-filtered seawater, placed in a water bath, and closed with lids. Each lid was fitted with a sampling port, an optical oxygen sensor (FireStingO_2_, PyroScience GmbH, Germany), and a magnetic stirrer. Before substrate addition, triplicate water samples for inorganic nutrient concentration quantification (ammonia, nitrite, nitrate, silicon) were taken through the sampling port at T0, filtered through 0.45 μm filters into 6 ml vials, and stored frozen (−20ºC; ammonia, nitrite, nitrate) or cold (4ºC; silicon) until analysis within <1 week. Furthermore, duplicate water samples for dissolved inorganic C (DIC) and ^13^C-DIC analysis were taken in 10 ml headspace vials, preserved by adding 10 μl saturated HgCl_2_ and stored at 4°C. For the substrate addition, 3.5 ml defrosted, homogenized labelled inactivated bacteria concentrate (equivalent to ∼80 mg org. C l^−1^; later on called “substrate bacteria”) was injected through the sampling port into the incubation chamber. Immediately afterwards (T1), water samples were taken for the analysis of ^13^C-DIC, DIC, and inorganic nutrient concentrations as described above, and the missing water in the incubation chamber was replaced with prefiltered seawater. Chambers were sealed off airtight with lids and oxygen consumption was measured continuously in the dark over a 12 h period at ambient temperature (∼12°C) with optical oxygen sensors. After 12 h (T2), water samples for DIC, ^13^C-DIC, and inorganic nutrient concentrations were taken again, and the incubations were terminated. Sponges were retrieved from incubation chambers (T3) and rinsed in pre-filtered seawater to remove any residuals of the labelled substrate. Individual specimens were photographed, and length, width, and height of sponges were measured with a ruler to the nearest millimeter. Sponges were scrapped off from any hard substrate they were attached to (e.g., oyster shells or stones) and the volume of each individual specimen was determined by water displacement. Subsequently, all sponges were weighed and preserved at −20ºC. Freshly sampled sponges served as background sample for natural abundance δ^13^C, δ^15^N, and δ^2^H-values and were kept frozen at −20ºC.

To quantify potential incorporation of ^2^H in dead sponge tissue (Fig. 2B), three sponge specimens were collected, transported to NIOZ-Yerseke, and gently washed with pre-filtered seawater from the Eastern Scheldt. Length, width, and height of the specimens were determined before they were scrapped off any hard substrate. Their tissue volume was measured via water displacement, and they were weighed. Each sponge was cut into equally sized pieces and kept in 4% formaldehyde solution for 10 min before the sponges were transferred to individual incubation chambers (V = 8.5 l). 85 ml ^2^H-water (equivalent to 1% ^2^H-enrichment) was added to each chamber that was filled subsequently with pre-filtered seawater. The chambers were closed with lids, placed in the water bath at ambient temperature, and incubated for 12 h. At the end of the incubation period, sponges were retrieved from the chambers and frozen (−20ºC).

To determine the contribution of sponges to oxygen consumption, DIC, ^13^C-DIC, and inorganic nutrient fluxes (Fig. 2C), incubation chambers (V = 1.4 l) were filled with pre-filtered seawater from the Eastern Scheldt and placed in a water bath that maintained ambient temperature. At T0, water samples for the quantification of DIC, ^13^C-DIC, and inorganic nutrients were taken as described above for the main treatment. Before water samples were taken again at T1, 0.56 ml defrosted and homogenized labelled substrate bacteria (equivalent to ∼80 mg org. C l^−1^) were injected through the sampling port in the lid into the incubation chamber. The incubation chambers were filled to the top to replace extracted water. They were closed airtight with a lid that was fitted with a sampling port, a magnetic stirrer, and an optical oxygen sensor, and oxygen consumption in the dark was recorded continuously for 12 h. At the end of the incubation period (T2), water samples for DIC, ^13^C-DIC, and inorganic nutrient concentrations were taken and preserved as described for the main treatment.

### Analysis of sponge tissue

Frozen sponge tissue (−20ºC) was freeze-dried and ground manually to fine powder. Ash-free dry mass (AFDM, g) was measured after oxidizing sub-samples (2 – 3 g) of freeze-dried sponges in a muffle furnace at 450°C for 4 h. Org. C content/ δ^13^C and TN content/ δ^15^N in sponge tissue were analyzed by packing ∼1 mg sponge powder in 8 × 5 mm pre-combusted (4 h at 450°C) silver capsules, acidifying them with 20 μL 2% HCl, and drying them at 60°C on a hot place. Afterwards, capsules were closed, and measured with a Flash EA 1112 elemental analyzer (Thermo Fisher Scientific, USA) coupled to a DELTA V Advantage Isotope Ratio Mass Spectrometer (Thermo Fisher Scientific, USA) via a ConFlo III interphase at NIOZ-EDS (EA-c-IRMS settings are presented in Supplementary Material A1). L-glutamic acid reference material USGS40 (δ^13^C_VPDB_: −26.2‰, δ^15^N_air_: −4.52‰) (US Geological Survey, Reston, Virginia, USA; (Qi et al., 2003)) and USGS41a (δ^13^C_VPDB_: +36.6‰, δ^15^N_air_: +47.6‰) (US Geological Survey, Reston, Virginia, USA; (Qi et al., 2016)) served as secondary reference material against which δ^13^C and δ^15^N data were normalized as described in (Sharp, 2017).

For the analysis of H content/ δ^2^H, ∼0.5–1.5 mg freeze-dried, powdered sponge tissue was packed in 5.25 × 3.2 mm silver capsules, closed, and measured with an EA IsoLink CN/OH elemental analyzer (Thermo Fisher Scientific, USA) coupled to a Flash IRMS Isotope Ratio Mass Spectrometer (Thermo Fisher Scientific, USA) at NIOZ-MMB (EA-TC-IRMS settings are presented in Supplementary Material A1). The mineral oil ‘NBS 22’ (No. 454, δ^2^H_VSMOW-SLAP_: −117±0.6‰; International Atomic Energy Agency IAEA, Vienna, Austria; (Schimmelmann et al., 2016)), Caribou Hoof Standard (CBS, δ^2^H_VSMOW-SLAP_: −197 ± −1.8‰; Environment Canada, Saskatoon, Saskatchewan, Canada) and Kudu Horn Standard (KHS, δ^2^H_VSMOW-SLAP_: −54.1±-0.6‰; Environment Canada, Saskatoon, Saskatchewan, Canada) served as secondary reference material for δ^2^H. δ^2^H data were normalized against CBS and KHS, before the δ^2^H values were corrected following Soto et al. (2017):

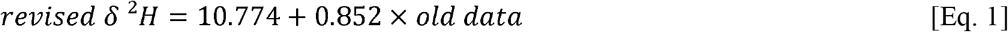

Stable isotope values were presented in δ notation relative to Vienna Pee Dee Belemnite (VPDB) for δ^13^C, relative to air for δ^15^N (Fry, 2006), and relative to Standard Mean Ocean Water (VSMOW) normalized to Standard Light Antarctic Precipitation (SLAP; VSMOW-SLAP) for δ^2^H (Coplen, 1995).

### Analysis of fatty acids in sponge tissue

Phospholipid-derived fatty acids (PLFAs) were extracted from ∼110–160 mg freeze-dried, powdered sponge tissue and ∼20 mg freeze-dried substrate bacteria following a modified Bligh and Dyer extraction method (Bligh & Dyer, 1959; Boschker, 2008). In the first step (Step A: Bligh and Dyer extraction), 15 ml methanol (HPLC grade, 99.8%), 7.5 ml chloroform (HPLC grade, 99.5%; used for sponge extraction) or 7.5 ml dichloromethane (DCM, HPCL grade; used for substrate bacteria extraction), and 6 ml MilliQ-water were mixed with the sponge powder and substrate bacteria powder, respectively, in pre-cleaned glass tubes (rinsed with HPLC grade methanol and hexane). The tubes were closed with Teflon coated lids and shaken on a shaker table for 2 h. Additional 7.5 ml chloroform or DCM was added to the tubes which were shaken shortly by hand before another 7.5 ml Milli-Q water was added. The tubes were stored at −20°C over night to let the solvent layers separate. The lower solvent layer containing the chloroform or DCM with the total lipid extract was transferred with a glass syringe to pre-weighted glass tubes. After the weight of the chloroform or DCM layer was determined, it was evaporated until complete dryness, 1 ml chloroform or DCM was added, and it was evaporated again. Subsequently, the total lipid extract was dissolved in 0.5 ml chloroform or DCM and in step 2 (Step B: Column chromatography) this extract was added to an activated silicic acid column (heated at 120ºC for 2 h; Silica gel 60, 0.063–0.200 mm, extra pure for column chromatography) to fractionate the total lipid extract into the different lipid classes. The total lipid extract was eluted on the column with 7 ml chloroform or DCM, 7 ml acetone (HPLC grade, 99.8%), and 15 ml methanol, and the methanol fraction containing the most polar lipid fraction was collected. This methanol fraction was evaporated and subsequently derivatized to fatty acid methyl esters (FAMEs) in step 3 (Step 3: Mild alkaline methanolysis): 1 ml methanol-toluene mix (1:1 volume/ volume), 20 μl of an internal standard (1 mg C19:0 FAME mL^−1^), and 1 ml methanolic NaOH (0.2 mol l^−1^) were added to the dried polar lipids and incubated for 15 min at 37°C. Subsequently, 2 ml *n*-hexane, 0.3 ml acetic acid (1 mol l^−1^), and 2 ml MilliQ-water were added, and the solution was mixed well. After the solvent layers separated, the (top) *n*-hexane layer was transferred to new glass tubes. Additional 2 ml *n*-hexane were added to the used glass tubes, the solvents in the tubes were mixed well, and the *n*-hexane layer was transferred again to the new tubes. A second internal standard (20 μl of 1 mg C12:0 FAME mL^−1^) was added and the *n*-hexane was evaporated to complete dryness. In a last step, the FAMEs were dissolved in 200 μl *n*-hexane and transferred to gas chromatograph (GC) glass vials.

C concentrations of individual FAMEs were measured with a Gas Chromatograph Trace GC Ultra (Thermo Fisher Scientific, USA) with a flame ionization detector (FID) using a polar BPX70 column (length: 50 m length, inner diameter: 0.32 mm, film thickness: 0.25 μm; SGE Analytical Science) (GC-FID settings are presented in Supplementary Material A2). Various reference material FAMEs (C12:0 FAME, C13:0 FAME, C14:0 FAME, C15:0 FAME, C16:0 FAME, C18:0 FAME, C19:0 FAME, C20:0 FAME, C22:0 FAME, C23:0 FAME, C24:0 FAME, C16:1ω7*t* FAME, C16:1ω7*c*, C18:1ω9*t*, 10Me-C16:0, 10Me-C18:0) from Schimmelmann Research (Indiana University Bloomington, USA) were combined in a mix and served as standards for quality control.

Individual peaks in the chromatograms were identified based on their equivalent chain length (ECL) in relation to ECLs of the internal standards C12:0 FAME and C19:0 FAME and the omnipresent peak of C16:0 FAME. For this, retention times (RTs) of peaks were converted to their ECLs as follows:

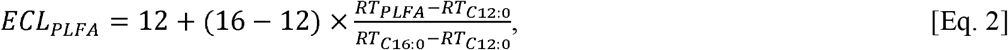

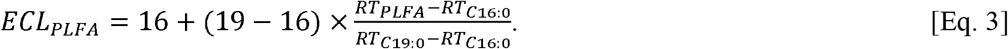

Equation 2 was used to calculate ECLs for PLFAs between C12:0 and C16:0, and equation 3 was used to estimate ECLs for PLFAs between C16:0 and C19:0. ECLs of PLFAs longer than C19:0 were calculated using equation 3.

ECLs were compared with a ECL reference list using the *R* (version 2.15.3) package *Rlims* (Soetaert, Provoost & van Rijswijk, 2015) (version 1.03). This ECL reference list was prepared based on RTs of certified FAME reference standards from Sigma-Aldrich (USA), Larodan (Sweden), Supelco® Analytical (USA), and from Schimmelmann Research that were measured on the same GC-FID with the same BPX70 column.

C concentrations of individual (identified) PLFAs (μg C g^−1^ DM sponge tissue or substrate bacteria) were calculated based on the peak area of the respective FAME (*A*_*FAME*_) in the chromatogram, the peak area of the internal standard C19:0 FAME (*A*_*19:0*_) in the chromatogram, and the concentration of the added internal standard (*c*_*19:0*_) using *Rlims* as follows:

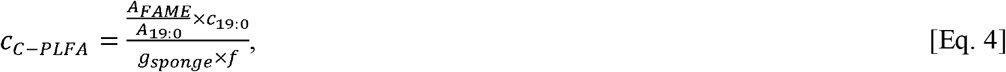

where *g*_*sponge*_ corresponds to the total amount of sponge tissue or substrate bacteria from which PLFAs were extracted and *f* is the fraction of chloroform or DCM that was recovered during the extraction.

H concentrations were subsequently calculated based on the C concentrations of the respective (identified) PLFAs and their chemical formula.

The δ^13^C_VPDB_ and δ^2^H_VSMOW-SLAP_ values (‰) of individual FAMES (technical replicates: 1 for C, 3 for H) were measured on a Delta V Advantage Isotope Ratio MS (Thermo Scientific, USA) coupled to a Trace 1310 GC (Thermo Scientific) with a Isolink II interface coupled to a Conflo IV (Thermo Scientific) using a BPX70 column (GC-IRMS settings are presented in Supplementary Material A3).

Individual peaks of the GC-IRMS chromatograms were identified based on their ECLs which were compared to a ECL reference list that had been prepared with the analytical standard PUFA No. 3 (Supelco® Analytical Products, USA), the certified reference material Supelco 37 Component FAME Mix (Supelco), and the internal standards C12:0 FAME and C19:0 FAME. Remaining unidentified peaks were identified by mass spectrometry (instrument series: 5977A MSD) coupled to a gas chromatograph (series: 7890B; Agilent Technologies, USA).

The isotopic values of the PLFAs were corrected for the addition of the methyl group using *Rlims*:

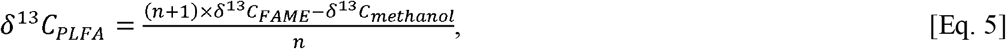

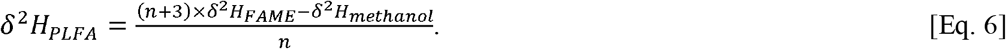

### Analysis of inorganic nutrients and dissolved inorganic carbon

Ammonia, nitrite, nitrate, and silicon concentrations (μmol l^−1^) were quantified with a SEAL QuAAtro analyzer (Bran+Luebbe, Germany), after warming the inorganic nutrient samples to room temperature for 24 h.

For the measurements of DIC concentrations (μmol l^−1^), He-gas was injected through the septum of the headspace vials and a ∼3 ml headspace was created. Excess sample was removed through a second syringe as described in (Moodley et al., 2000) and ∼2.5 ml of this excess water was used to quantify the DIC concentration on an Apollo SciTech DIC analyzer (AS-C3; Apollo SciTech, USA). All inorganic DIC in the headspace vial was transformed to gaseous CO_2_ by acidification with 10 μl concentrated H_3_PO_4_ per 1 ml water. A 500 μl gas sample was taken from the headspace and injected into a Thermo Delta V continuous flow IRMS to measure δ^13^C_VPDB_ (‰) of the CO_2_ (Gillikin & Bouillon, 2007).

## Calculations

The sponge condition index *CI* (−) was calculated as described in (Lüskow et al., 2019):

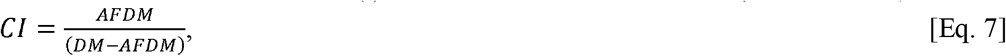

The incorporation (*I*) of tracer C (μmol tracer C mmol C_sponge_^−1^), N (μmol tracer N mmol C_sponge_ ^−1^), and H (μmol tracer H mmol C_sponge_ ^−1^) into sponges was calculated as described in (Stratmann et al., 2023):

The ratios of ^13^C/ ^12^C, ^15^N/ ^14^N, and ^2^H/ ^1^H in the bulk sponge material and PLFAs, i.e., *R*_*bulk*_ and *R*_*PLFA*_, is

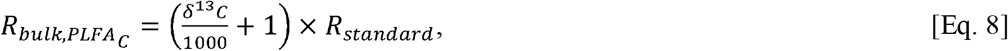

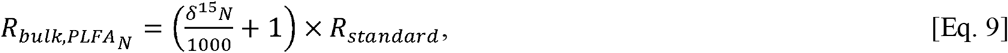

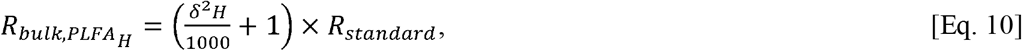

where *R*_*standard*_ for C is 0.01118, for N is 0.0036765, and for H it is 0.00015576 (Fry, 2006). The fraction (*F*) of the heavy isotopes in the bulk sponge material, bacterial substrate, and PLFAs was calculated as:

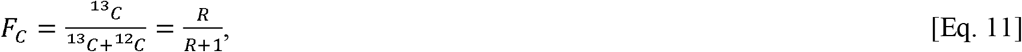

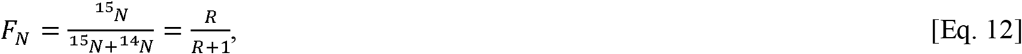

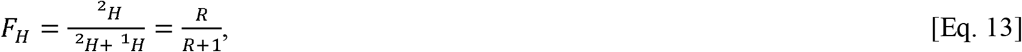

The incorporation of tracer C, N, and H into the sponge tissue is:

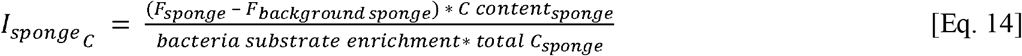

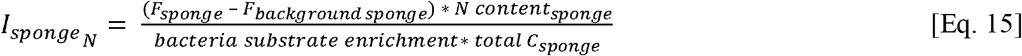

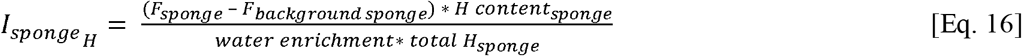

In this study, substrate bacteria ^13^C enrichment was 28.6 at%, substrate bacteria ^15^N enrichment was 51.0 at%, and water enrichment was 1 at%.

The incorporation of ^13^C (*I*_*C*_) and ^2^H (*I*_*H*_) into PLFAs is:

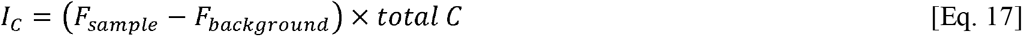

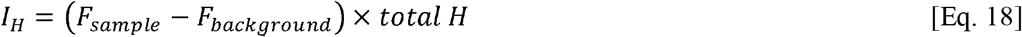

For the calculations of ^13^C and ^2^H incorporation into PLFAs, only PLFAs were considered that were detected in at least 3 samples per treatment.

Abiotic incorporation of ^2^H into PLFAs was corrected as follows:

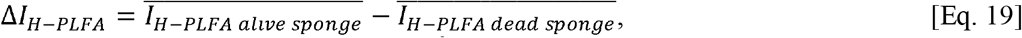

where 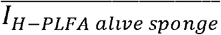 is the mean ^2^H incorporation into an individual PLFA of alive sponges and 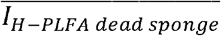 is the mean ^2^H incorporation into a PLFA of dead sponges.

Blank-corrected and biomass-corrected fluxes (*Flux*) of inorganic nutrients (nitrate, nitrite, ammonia, silicon) (μmol mmol C^−1^ d^−1^) were calculated as described in (Stratmann et al., 2024):

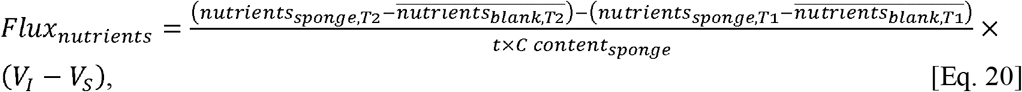

*nutrients*_*sponge*_ is the inorganic nutrients concentration (μmol l^−1^) in the incubation chamber with a sponge and 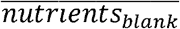 is the mean concentration of inorganic nutrients in the incubation chamber without a sponge (*n* = 3). Sample *T1* was taken at the beginning of the incubation after adding labelled substrate bacteria, and *T2* corresponds to the end of the incubation after 12 h. *t* is the incubation time (=0.5 d) and *C content*_*sponge*_ is the org. C content of the sponge (mmol C). *V*_*I*_ is the volume of the incubation chamber (= 8.5 l) and *V*_*S*_ is the volume of the sponge (l).

Blank-corrected and biomass-corrected fluxes of DIC (μmol DIC mmol C^−1^ d^−1^) were calculated as described for inorganic nutrients.

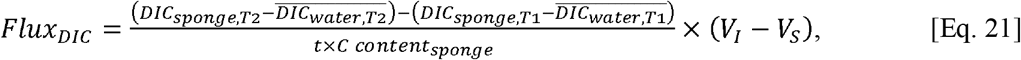

Blank-corrected and biomass-corrected fluxes of tracer-C DIC (μmol tracer-DIC mmol C^−1^ d^−1^) was calculated by combining Eq. 14 and Eq. 20:

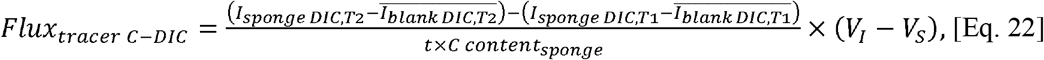

where *I*_*sponge DIC*_ is the production of tracer C-DIC in the incubation chamber with a sponge and *I*_*water DIC*_ is the production of tracer C-DIC in the incubation chamber without a sponge.

Blank-corrected and biomass-corrected fluxes of O_2_ (mmol O_2_ mmol C^−1^ d^−1^) were calculated as follows:

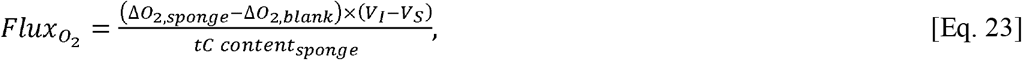

where ΔO_2_ (mmol O_2_ l^−1^ d^−1^) is the average decrease in oxygen over time. This decrease was calculated by linear regression, after air saturation (%) measured with the oxygen meter was converted to absolute oxygen concentration (μmol O_2_ l^−1^) following (Weiss, 1970) using the R package *marelac* (Soetaert, Petzoldt & Meysman, 2010).

### Statistical analysis

To test whether oxygen, inorganic nutrient, DIC, and ^13^C-DIC fluxes were significantly different from 0 mmol mmol C^−1^ d^−1^ or 0 μmol mmol C^−1^ d^−1^, respectively, 1-sided Student’s t-tests (α = 0.05) were applied in *R* (version 4.3.0) to normally distributed data, after normality was confirmed by the Shapiro-Wilk normality test. Non-normally distributed data were tested by a 1-sample Wilcoxon test (α = 0.05).

Differences in uptake rates of ^2^H by alive sponges and dead sponges were tested by the non-parametric Kruskal-Wallis rank sum test *H* (α = 0.05), after the Bartlett test of homogeneity of variances (α = 0.05) revealed that the variances of the two groups were different.

All data are expressed as mean ± standard error (SE).

## Results

### Biogeochemical characteristics of *H. panicea*

Sponges had a volume of 76.4 ± 14.1 ml, a wet mass (WM) of 114 ± 21.8 g, and a water content of 14.0 ± 11.0% WM. The dried sponge tissue consisted of 12.8 ± 1.29% org. C, 2.90 ± 0.31% TN, and 2.47 ± 0.30% H. The mean δ^13^C-value of the sponge tissue was −18.6 ± 0.03‰, the mean δ^15^N-value was 8.66 ± 0.02‰, and the mean δ^2^H-value was −74.6 ± 1.64.‰. Sponge condition index *CI* was 0.38 ± 0.08.

Sponges contained mostly long-chain fatty acids (LCFA, i.e., fatty acid with ≥24 C atoms; 62.1 ± 2.40%), saturated fatty acids (SFA; 22.8 ± 1.55%), and monosaturated fatty acids (MUFA; 11.1 ± 0.68%). Most analyzed PLFAs were sponge-specific (56.6 ± 2.78%) and bacteria-specific PLFAs (15.8 ± 0.98%), or they belonged to the category ‘others’ (27.6 ± 1.86%).

Natural abundance stable C isotope values (δ^13^C) of investigated PLFAs ranged from −29.7‰ (5^th^ percentile) to −15.6‰ (95^th^ percentile) (Fig. 3, upper panel) and natural abundance stable H isotope values (δ^2^H) of studied PLFAs were between −348‰ (5^th^ percentile) and −56.3‰ (95^th^ percentile) (Fig. 3, lower panel). The PLFA which was most depleted in δ^13^C was C20:1ω9c (−30.6 ± 0.71‰) and the PLFA which was most depleted in δ^2^H was C14:0 (−368 ± 18.7). In comparison, the PLFA that was most enriched in δ^13^C was C16:1ω7c (−9.62 ± 2.41‰) and the PLFA that was most enriched in δ^2^H was C15:1ω5c (−64.9 ± 5.48). Bacteria-specific PLFAs were more enriched in δ^13^C (mean ± SE: −17.4 ± 0.84‰; median: −17.6‰; 5^th^ – 95^th^ percentile range: −23.3‰ – −8.67‰) than sponge-specific PLFAs (−22.4 ± 1.12‰; −21.9‰; −32.7‰ – −18.3‰), whereas bacteria-specific PLFAs were more depleted in δ^2^H (−128 ± 10.8; −110‰; −322‰ – −52.0‰) than the sponge-specific PLFAs (−153 ± 5.49; −142‰; −236‰ – −96.3‰).

**Figure 3.**
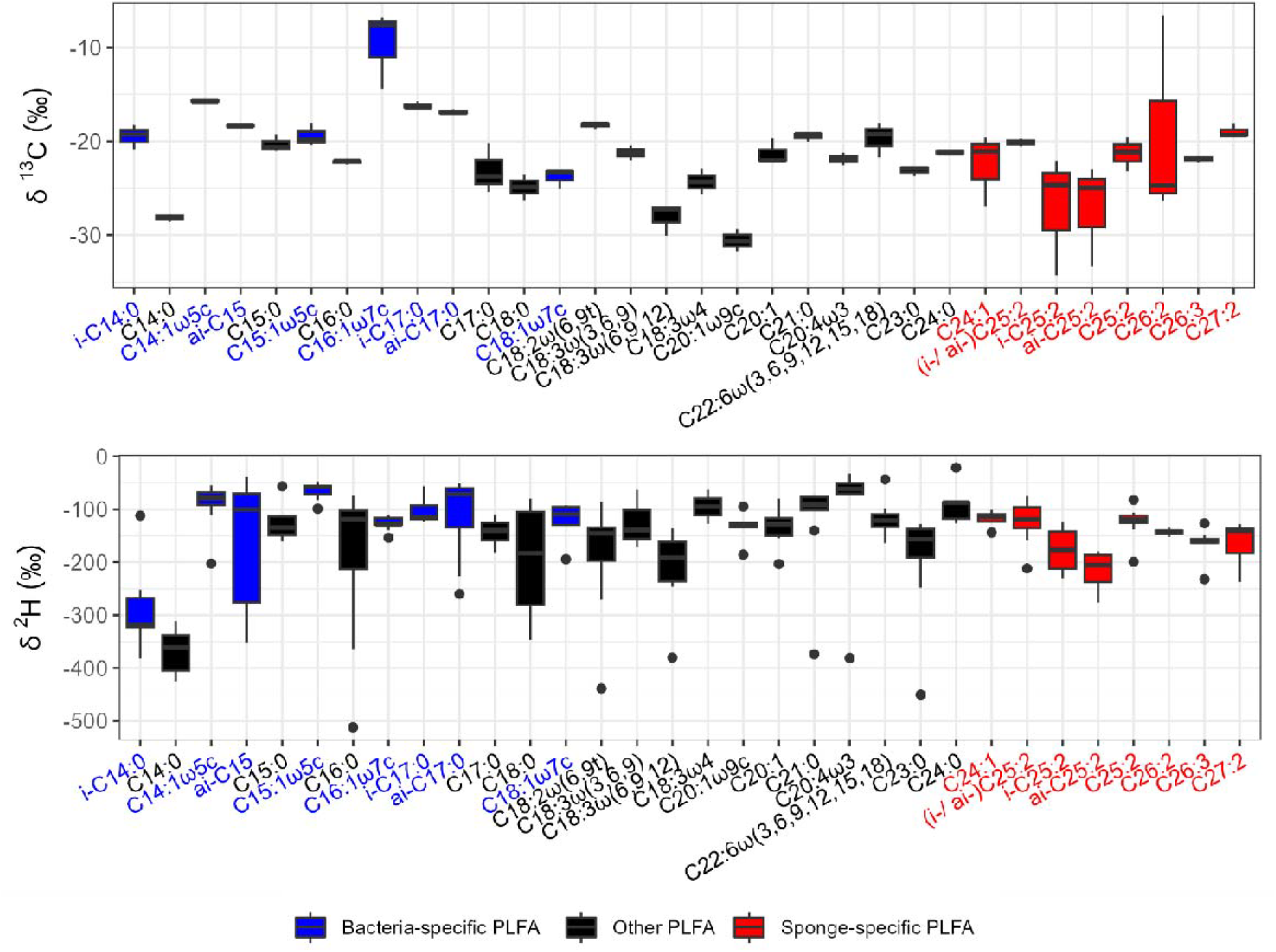
Natural abundance δ^13^C_VPDB_ (upper panel) and δ^2^H_VSMOW-SLAP_ (lower panel) composition (‰) of phospholipid-derived fatty acids in *Halichondria* (*Halichondria*) *panicea*.

### Oxygen, dissolved inorganic carbon, and inorganic nutrient fluxes

In the incubation experiment, alive sponges consumed oxygen (0.72 ± 0.59 mmol O_2_ mmol C^−1^ d^−1^) and released ^13^C-DIC (0.29 ± 0.04 mmol mmol C^−1^ d^−1^) and ammonia (124 ± 11.2 μmol mmol C^−1^ d^−1^). At the same time, they took up nitrite (4.66 ± 1.04 μmol mmol C^−1^ d^−1^), nitrate (8.31 ± 5.12 μmol mmol C^−1^ d^−1^), and silicon (33.7 ± 2.66 μmol mmol C^−1^ d^−1^). However, only the ammonia, nitrite, silicon, and ^13^C-DIC fluxes were statistically significantly different from 0 μmol mmol C^−1^ d^−1^ (Table 1).

**Table 1.** Results of 1-sided Student’s t-tests (α = 0.05) and 1-sample Wilcoxon tests (α = 0.05) to assess whether oxygen consumption rates (mmol O_2_ g mmol C^−1^ d^−1^), ^13^C-DIC and inorganic nutrient fluxes (μmol mmol C^−1^ d^−1^) were significantly different from 0 (H_0_: μ = 0, H_1_: μ ≠ 0). Symbols: \**p*-value≤0.05, □ *p*-value≤0.01

| Flux type | t-value | df | p-value | Mean | 95% Confidence interval |
| --- | --- | --- | --- | --- | --- |
| O<sub>2</sub> | -2.47 | 3 | 0.0904 | -0.72 | -1.65, 0.21 |
| NH<sub>4</sub><sup>+</sup> | 11.1 | 4 | 0.0004□ | 124 | 93.4, 155 |
| NO<sub>2</sub><sup>-</sup> | -4.49 | 4 | 0.0109□ | -4.66 | -7.54, -1.77 |
| NO<sub>3</sub><sup>-</sup> | -1.62 | 4 | 0.1801 | -8.31 | -22.5, 5.92 |
| Si | -12.6 | 4 | 0.0002□ | -33.7 | -41.1, -26.3 |
| <sup>13</sup>C-DIC | 8.23 | 4 | 0.0012□ | 292 | 193, 390 |

### Total C, N, and H incorporation

Alive sponges incorporated 17.0 ± 4.47 μmol tracer C mmol C^−1^ d^−1^ (corresponding to 4.82 ± 1.27 μmol ^13^C mmol C^−1^ d^−1^), 3.56 ± 0.93 μmol tracer N mmol C^−1^ d^−1^ (corresponding to 2.46 ± 1.16 μmol ^15^N mmol C^−1^ d^−1^), and 49.1 ± 19.7 μmol tracer H mmol C^−1^ d^−1^ (corresponding to 0.49 ± 0.20 μmol ^2^H mmol C^−1^ d^−1^). This ^2^H incorporation was significantly higher than the incorporation of tracer H/ ^2^H in dead sponges (19.2 ± 6.20 μmol tracer H mmol C^−1^ d^−1^, corresponding to 0.19 ± 0.06 μmol ^2^H mmol C^−1^ d^−1^; *H*(1) = 5, *p* = 0.03).

### C and H incorporation into fatty acids

Substrate bacteria contained ^13^C-enriched PLFAs with a total value of 4.09 ± 0.36 mg ^13^C g^−1^ DM bacteria; PLFAs with the highest ^13^C-enrichment were C16:0, C16:1ω7c, C16:1ω9, C18:0, and C18:1ω7c (Fig. S1).

The consumption of δ^13^C-enriched substrate bacteria by alive *H. panicea* resulted in a strong enrichment in δ^13^C of the PLFAs i-C14:0 (mean ± SE: 2021 ± 512; median: 2798‰; 5^th^ – 95^th^ percentile range: 634‰ – 2909‰), C16:0 (2130 ± 344‰; 2369‰; 1136‰ – 2873‰), C16:1ω7c (7802 ± 1316‰; 9842‰; 4225‰ – 10018‰), i-C17:0 (800 ± 308‰; 1009‰; −134‰ – 1330‰), C18:2ω(2,9t) (1680 ± 380‰; 1507‰; 1035‰ – 2567‰), and C22:6ω(3,6,9,12,15,18) (1219 ± 339‰; 1489‰; 639‰ – 1610‰) (Fig. 4, upper panel). The incubation of these alive sponges in δ^2^H-enriched water led to a δ^2^H enrichment of most PLFAs (Fig. 4, lower panel; Table 1) which ranged from −207 to 646‰ (5^th^ – 95^th^ percentile range; mean ± SE: 104 ± 14.2‰, median: 50.8‰). The three PLFAs that were most enriched in δ^2^H were C18:2ω(6,9t) (864 ± 1316‰; 41.1‰; 772‰ – 923‰), C18:1ω7c (297 ± 94.3‰; 298‰; 166‰ – 402‰), and C24:1 (519 ± 186‰; 293‰; 200‰ – 1063‰), whereas the three PLFAs with the least δ^2^H-enrichment were C14:0 (−285 ± 13.3‰; −278‰; −316‰ – −247‰), ai-C25:2 (−91.6 ± 29.8‰; −103‰; −132‰ – −61.3‰), and C23:0 (−7.24 ± 81.2‰; −98.1‰; −101‰ – −54.9‰). In comparison, dead sponges (Fig. S2) had δ^2^H values of −134 ± 4.56‰ (median: −123‰; 5^th^ – 95^th^ percentile range: −272 – −54.7‰). When δ^2^H values of the individual PLFAs were compared, the 5^th^ to 95^th^ percentile ranges of PLFAs from dead-sponge incubations vs. alive sponge incubations did not overlap in 56.3% of the analyzed PLFAs (Table S2). In 84.4% of the studied PLFAs, the 25^th^ to 75^th^ percentile ranges of δ^2^H values did not overlap.

**Figure 4.**
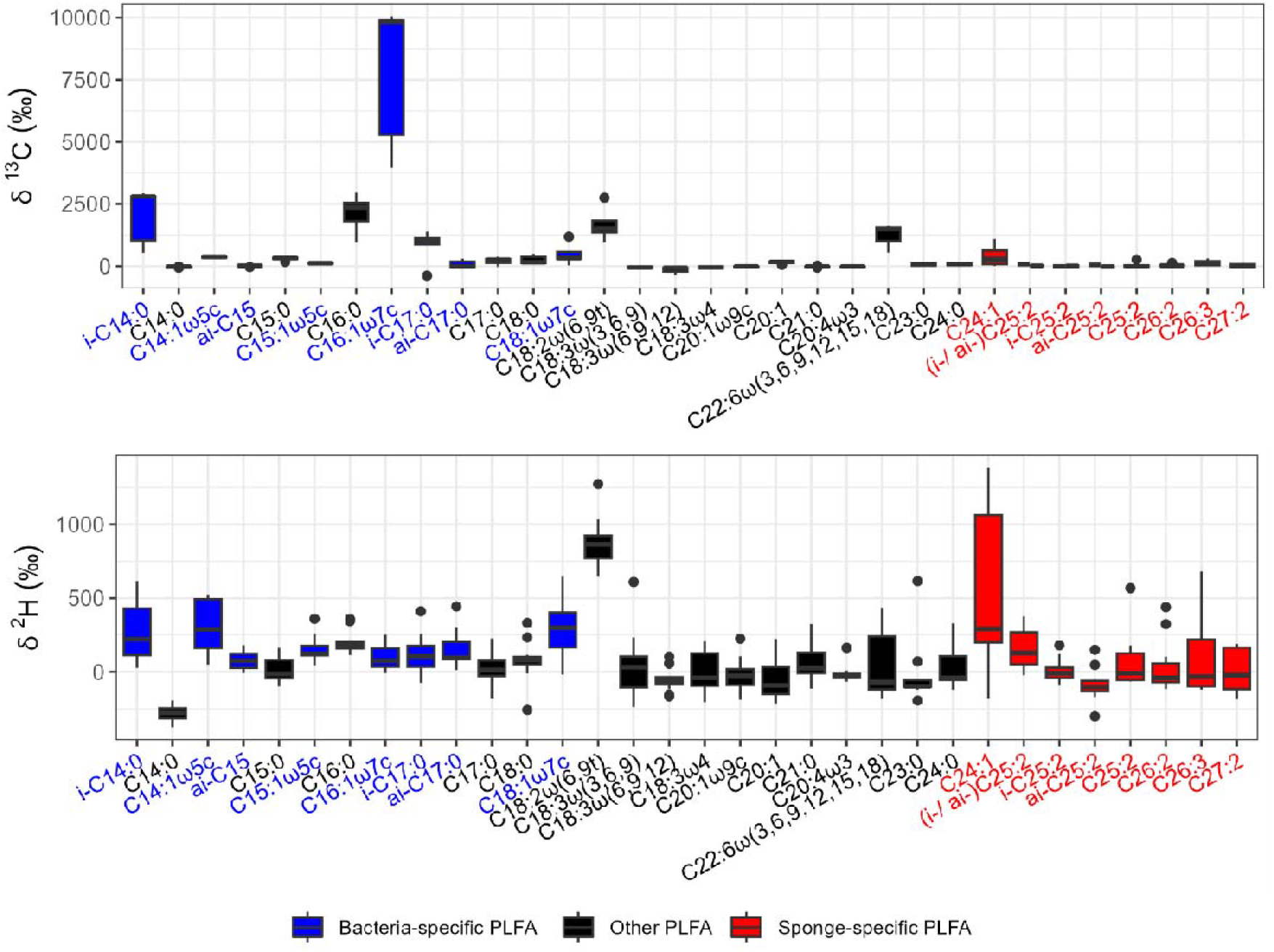
Stable isotopic enrichment of PLFAs extracted from *H. panicea* specimens incubated for 12 h in ^2^H-enriched water in the presence of ^13^C-enriched substrate bacteria. (Upper panel) δ^13^C_VPDB_ values (‰) of PLFAs and (lower panel) δ^2^H_VSMOW-SLAP_ values (‰) of PLFAs.

The investigated PLFAs contained 5.61 ± 1.81 μg ^13^C g^−1^ DM sponge tissue, of which 51.7% (2.90 ± 1.01 μg ^13^C g^−1^ DM sponge tissue) was present in C16:1ω7c and 23.2% (1.30 ± 0.47 μg ^13^C g^−1^ DM sponge tissue) was present in C16:0 (Fig. 5). Bacteria-specific PLFAs deriving from substrate bacteria and from the sponge microbiome consisted of 3.38 ± 1.11 μg ^13^C g^−1^ DM sponge tissue, whereas sponge-specific PLFAs had 0.57 ± 0.20 μg ^13^C g^−1^ DM sponge tissue, and 1.67 ± 0.55 μg ^13^C g^−1^ DM sponge tissue was built in the other PLFAs.

**Figure 5.**
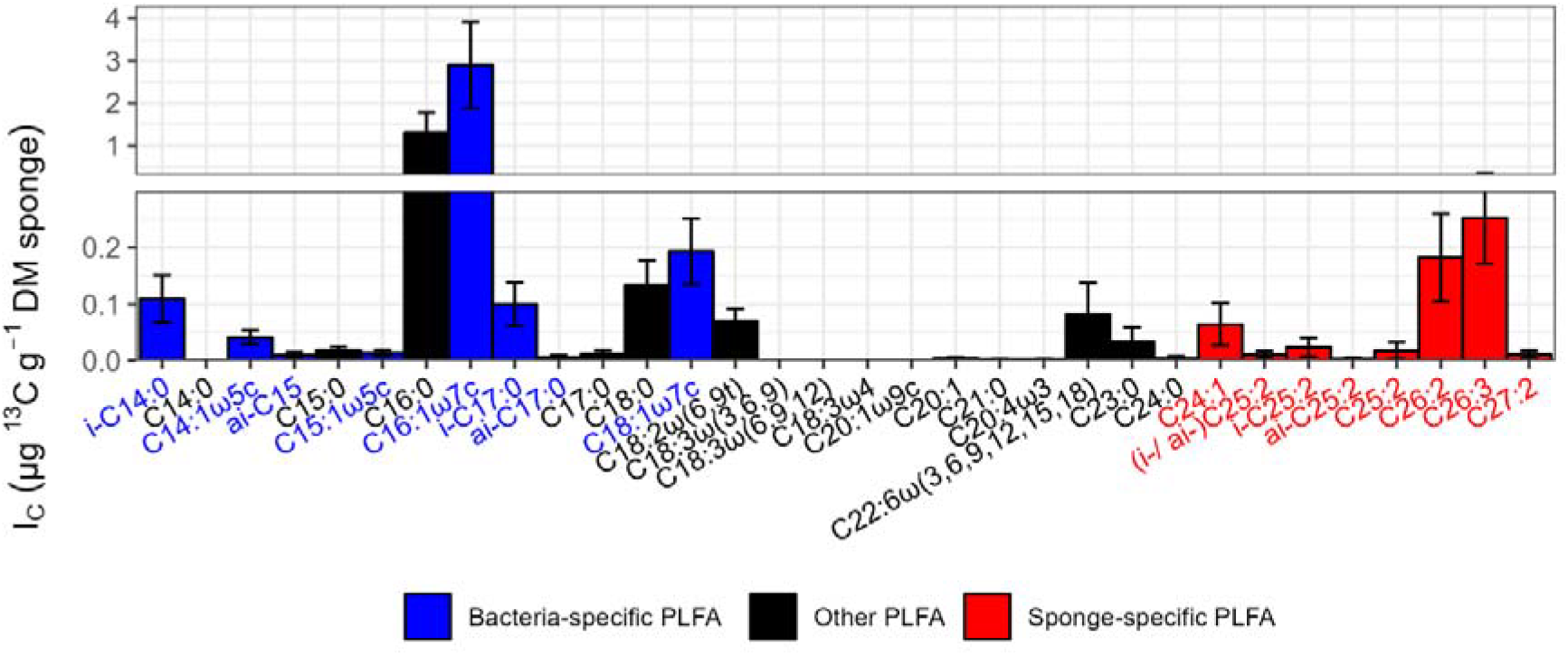
Incorporation of ^13^C (I_C_) into phospholipid-derived fatty acids (PLFA) (μg ^13^C g^−1^ DM sponge) from alive sponges that were fed with a ^13^C-enriched substrate bacteria. Error bars represent 1 SE.

PLFAs of alive sponges included 5.43 ± 1.42 ng ^2^H g^−1^ DM sponge tissue, of which 20.9% were present in C26:2 (1.13 ± 0.36 ng ^2^H g^−1^ DM sponge tissue), 15.6% in C26:3 (0.85 ± 0.33 ng ^2^H g^−1^ DM sponge tissue), and 10.9% in C16:0 (0.59 ± 0.15 ng ^2^H g^−1^ DM sponge tissue) (Fig. S3, upper panel). Most ^2^H was incorporated into sponge-specific PLFAs (2.64 ± 0.83 ng ^2^H g^−1^ DM sponge tissue), followed by bacteria-specific PLFAs (1.14 ± 0.30 ng ^2^H g^−1^ DM sponge tissue) and other PLFAs (1.6582 ± 0.44 ng ^2^H g^−1^ DM sponge tissue).

The amount of ^2^H present in PLFAs from dead sponge tissue was about one order of magnitude lower than the amount of ^2^H present in PLFAs from alive sponges and accounted for 0.56 ± 0.06 ng ^2^H g^−1^ DM sponge tissue (Fig. S3, lower panel). Hence, the ^2^H uptake in individual PLFAs after correcting for the abiotic incorporation of ^2^H (Δ*I*_*H-PLFA*_) ranged from in total 0.98 ng ^2^H g^−1^ DM sponge tissue for all bacteria-specific PLFAs to in total 1.29 ng ^2^H g^−1^ DM sponge tissue for all other PLFA, and to in total 2.59 ng ^2^H g^−1^ DM sponge tissue for all sponge-specific PLFAs (Fig. 6).

**Figure 6.**
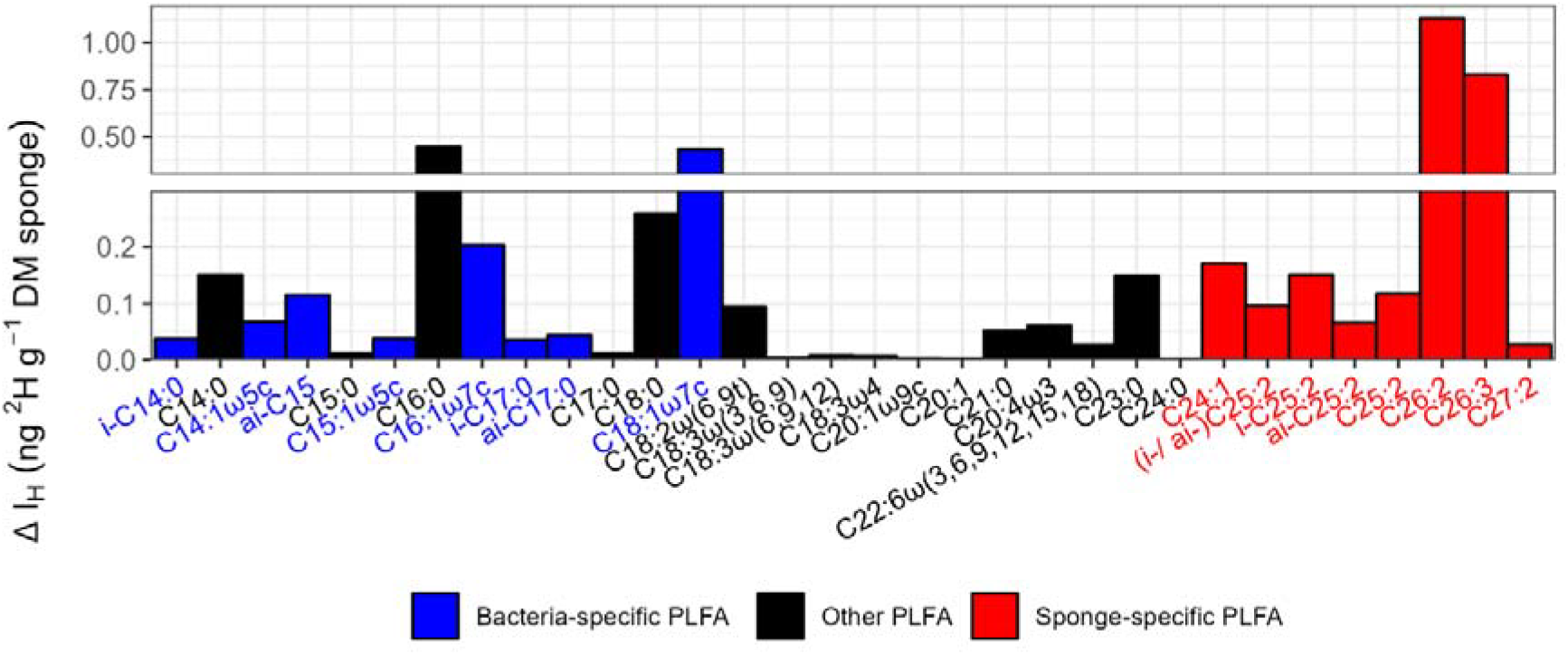
Incorporation of ^2^H into phospholipid-derived fatty acids (PLFA) after correcting for abiotic 2H incorporation (Δ*I*_*H-PLFA*_) (ng ^2^H g^−1^ dry mass sponge) from incubations of alive sponges.

## Discussion

Here, we measured oxygen consumption and inorganic nutrient fluxes in *H. panicea* from the Eastern Scheldt (Netherlands) to better understand its ecology. Furthermore, we studied the uptake of substrate bacteria by the sponge via assessing the incorporation of ^13^C and ^15^N into bulk sponge tissue, and the incorporation of ^13^C into PLFAs. Measurements of ^2^H uptake in bulk sponge tissue and in PLFAs were used to evaluate whether deuterated water can be used to determine metabolic activity of individual specimens.

### Ecology of *H. panicea* in the Eastern Scheldt

*H. panicea* is one of 18 sponge species present in the Eastern Scheldt (Van Soest et al., 2007; Bos et al., 2016). There, it grows as epibiont on macroalgae and on *Magallana gigas* (Thunberg, 1793) shells, but also on roof tiles and clinker bricks (Stratmann, personal observations), which originated from a drowned medieval village (clinker bricks) or were spread on the sediment in the last century to serve as hard substrate for shellfish larvae (roof tiles). In the Kiel Bight (Germany, Baltic Sea), in comparison, *H. panicea* uses mostly red macroalgae as substrate (Barthel, 1986, 1988).

In the Baltic Sea, *H. panicea*’s condition index *CI* ranged from ∼0.38 in February 1985 (winter) to 0.9 in June/ July 1984 (summer) in the Kiel Bight (Barthel, 1986) and from 0.53 in February 2016 (winter) to 1.14 in May/ June 2016 (spring/ summer) in Kerteminde Fjord (Denmark) (Lüskow et al., 2019). In contrast, in the Eastern Scheldt, *H. panicea*’s *CI* was 0.38 in August 2019 (summer) and therefore similar to the CI value of *H. panicea* from the Kiel Bight in winter. This is very surprising, but potentially related to the specific environmental conditions in the Eastern Scheldt. Primary production and chlorophyll a-concentration as a proxy for the phytoplankton community have decreased since 1995 (Smaal et al., 2013; Horn et al., 2023), after the construction of the storm surge barrier in the 1980s had resulted in an increase in chlorophyll a concentration (Wetsteyn & Kromkamp, 1994). This decrease in phytoplankton was caused by an excess cultivation of bivalves in the bay (Smaal et al., 2013; Jiang et al., 2020) that may be exposed to overexploitation if the shellfish stocks expand further (Smaal et al., 2013). Bacterioplankton as other food source for *H. panicea* was last studied in the Eastern Scheldt in 1984 during the construction of the storm surge barrier (Laanbroek et al., 1985) and we lack more recent data to assess whether its density and biomass increased or decreased over time. However, we except a trend comparable to the trend of phytoplankton because mussels (Hollibaugh, Fuhrman & Azam, 1980; Prieur, 1981; Birkbeck & McHenery, 1982) and oysters ((Douillet, 1993); Stratmann, unpublished data) also filter bacterioplankton out of the water column. Therefore, the low *CI* of *H. panicea* is an indication that this sponge species is disadvantaged compared to the bivalves and likely starving.

Potential starvation of *H. panicea* could also be assessed by investigating their silicon uptake. (Frøhlich & Barthel, 1997) studied silicon uptake rates of *H. panicea* in well-fed specimens, starved specimens, and re-fed specimens at 15°C. During starvation, the authors measured uptake rates that were 16% lower than the uptake rates of well-fed specimens (well fed specimens: 1.01 μmol Si g AFDM^−1^ h^−1^, starving specimens: 0.16 μmol Si g AFDM^−1^ h^−1^). Re-fed specimens had a silicon uptake of 0.33 μmol Si g AFDM^−1^ h^−1^. In this study, we measured a silicon uptake rate of 1.86 ± 0.51 μmol Si g AFDM^−1^ h^−1^ which corresponds more to a silicon uptake rate of well-fed instead of re-fed specimens. Hence, our data are inconclusive and we cannot be sure whether *H. panicea* in the Eastern Scheldt is indeed starving or not.

However, feeding org. C-rich inactivated substrate bacteria to potentially starving *H. panicea* specimens strongly affected their oxygen consumption rate: The oxygen consumption rate of specimens from this study (i.e., 219 ± 70.8 μmol O_2_ g DM sponge^−1^ h^−1^ or 807 ± 129 μmol O_2_ g AFDM sponge^−1^ h^−1^) was one to two orders of magnitude higher than the oxygen consumption rates of non-fed specimens that (Barthel, 1988; Kumala & Canfield, 2018; Kumala, Thomsen & Canfield, 2023) studied. This difference is large, but can be explained by the discrepancy of org. C content in the food source *H. panicea* usually encounter in its natural environment and the amount of C available in the ^13^C-enriched substrate bacteria that was fed in this experiment. In nature, *H. panicea* can maintain its metabolic activity when it filters phytoplankton with an org. C content of minimum 0.03 mg org. C l^−1^ or heterotrophic bacterioplankton and cyanobacteria with an org. C content of minimum 0.05 – 0.10 mg org. C l^−1^ (Riisgård, Kumala & Charitonidou, 2016). In this experiment, however, the sponges were fed with a substrate containing ∼80 mg org. C l^−1^, i.e., an org. C dose three orders of magnitude higher than they require for metabolic maintenance. So, *H. panicea* and its microbiome reacted to the availability of a very large, nutrient-rich food pool by increased respiration.

### Fatty acid composition of *H. panicea* and potential biosynthetic pathways

*H. panicea* and its microbiome incorporated ^13^C and ^2^H into bacteria-specific, sponge-specific, and unspecific (or other) PLFAs. Whereas ^13^C originated from the ^13^C-enriched substrate bacteria the sponge were fed, the source of ^2^H was the ^2^H-enriched ambient seawater in which the sponges were incubated for 12 h. This water is the dominant H source for the biosynthesis of fatty acids (Baillif et al., 2009). In fact, at even CH_2_ sites all H originate exclusively from water, whereas at odd CH_2_ sites, water is still the main H source, but non-exchangeable H from glucose can be also be incorporated (Baillif et al., 2009). Hence, by comparing ^13^C-enriched PLFAs from the (dead) substrate bacteria with ^13^C-enriched PLFAs from the sponge and its associated bacteria, and subsequently linking it with information about ^2^H-enriched PLFAs from the sponge and its microbiome, we can better constrain fatty acid biosynthesis: When a specific PLFA from *H. panicea* was enriched in ^2^H, this PLFA was actively produced via elongation or desaturation. When a specific PLFA was enriched in ^13^C, but not in ^2^H, this PLFA was directly taken up from the substrate bacteria and not produced via any ^13^C-enriched PLFA precursor. In comparison, when a PLFA was enriched in ^13^C and ^2^H, it was actively produced during the course of the experiment, but a fraction might have been taken up from the substrate bacteria, too, if it was present in it. If it was not present in the substrate bacteria, it would have been exclusively build by elongation and desaturation of ^13^C-enriched PLFA precursors present in the substrate bacteria. When a PLFA from the sponge was neither enriched in ^13^C nor in ^2^H, it originated from another food source and the sponge or its microbiome could not produce it themselves.

Living *H. panicea* contained several ^13^C-enriched bacteria-specific PLFAs, such as *i*-C14:0, *ai*-C15:0, *i*-C17:0, and *ai*-C17:0 (Fig. 5), though they were not present in the substrate bacteria (Fig. S1). As theses PLFAs were also enriched in ^2^H (Fig. 6), they were likely biosynthesized by the microbiome of *H. panicea* using ^13^C-enriched PLFA precursors from the substrate bacteria.

Sponge-specific LCFA were biosynthesized via a sequence of elongation and desaturation steps by intrinsic elongases and desaturases (encoded in the sponge genome) (The UniProt Consortium, 2023) using bacterial shorter-chain PLFAs derived from the substrate bacteria or from the sponge microbiome as precursors. For instance, the ^13^C- and ^2^H-enriched C24:1 [likely C24:1ω(5,9) (Ando et al., 1998; Rod’kina, Latyshev & Imbs, 2003)] might originate from C16:0 which underwent four rounds of elongation through C2 extension, followed by desaturation (Monroig & Kabeya, 2018), as Koopmans et al., (2015) had confirmed the bacterial origin of this PLFA via ^13^C labelling experiments. The branched *ai*-C25:2 [likely *ai*-C25:2ω(5,9) (Ando et al., 1998; Rod’kina, Latyshev & Imbs, 2003)] was derived from non- ^13^C-enriched bacterial *ai*-C15:0 (or *ai*-C17:0) from the sponge microbiome and underwent sequential elongation to C25:0 before it was desaturated at the position (5, 9) by sponge desaturases. The longer C26 PLFAs might originate from elongations of C14:0 (Hahn et al., 1988), which was present in the substrate bacteria (Fig. S1), and subsequently desaturated to from C26:2 [likely C26:2ω(5,9) (Ando et al., 1998; Rod’kina, Latyshev & Imbs, 2003)] and C26:3 [likely C26:3ω(5,9,19) (Ando et al., 1998; Rod’kina, Latyshev & Imbs, 2003)]. Alternatively, these PLFAs might derive from C16:0 with one elongation step less.

In summary, our findings provide evidence that the sponge microbiome biosynthesizes bacterial-specific PLFAs that provide the initial structural framework which is subsequently transformed by sponge enzymes into the long-chain derivatives.

### Deuterated water to measure metabolic activity in *H. panicea*

Measuring the faunal uptake of substrate-specific stable isotopes (^13^C, ^15^N) provides high-quality results for feeding activity when the target animals have been studied in detail and extensive knowledge exists about their feeding types and diet preferences. However, it is challenging to apply the traditional dual stable isotope approach to measure activity of less well studied organisms, like deep-sea fauna. Deep-sea benthos rarely encounters the type of substrate we may provide them (e.g. fresh phytodetritus or C-rich bacteria; but (Billett, Lampitt & Rice, 1983)) and knowledge about the natural diet of most deep-sea taxa is limited (e.g., (Ingels et al., 2010; Guilini et al., 2010; Pape et al., 2013)). Additionally, providing excessive amounts of food in one pulse to food-limited ecosystems may lead to wrong assumptions about ecosystem functioning as the organisms could react with increased metabolic activities compared to their activity during periods of starvation (Larsson, Lundälv & Van Oevelen, 2013; Maier et al., 2019).

In comparison, applying the substrate-independent stable isotope ^2^H (also called “substrate-agnostic isotope tracer” (Callaghan et al., 2023)) present in deuterated water provides information about the metabolic activity of an organism independent of its feeding activity. In fact, combined with substrate-specific stable isotopes, it allows to gain simultaneous insight in the transfer of C and N in the food web and about the metabolic activity of its individual members.

In this pilot study, living *H. panicea* incorporated 0.49 ± 0.20 μmol ^2^H mmol C^−1^ d^−1^ (equivalent to 0.55 ± 0.06 μmol ^2^H g WM sponge^−1^ d^−1^, 3.86 ± 0.86 μmol ^2^H g DM sponge^−1^ d^−1^, or 14.2 ± 1.68 μmol ^2^H g AFDM sponge^−1^ d^−1^) while feeding on the ^13^C- and ^15^N-enriched substrate bacteria. This resulted in bulk stable isotope uptake ratios of 8.67 ± 0.07 (^13^C/^15^N-ratio), 0.57 ± 0.12 (^13^C/^2^H-ratio), and 6.52×10^−2^ ± 1.35×10^−2^ (^15^N/^2^H-ratio). Hence, C and N transfer can be measured simultaneously with metabolic activity and even information about biosynthetic pathways of fatty acid production can be gained. However, strengths and weaknesses of this approach need further testing: For instance, the metabolic activity of fauna with different biological traits should be measured in the presence and absence of a food source. Other tests should include the metabolic activity of fauna in different growth stages and specimens of the same species should be exposed to different ambient temperatures. In addition, we recommend to focus more on compound-specific uptake rates of ^2^H (e.g., (Pilecky et al., 2021, 2022)) because 40% of the ^2^H measured as bulk uptake was H exchange in noncovalently bound H (Schimmelmann et al., 2001; Sessions et al., 2004; Lis, Schimmelmann & Mastalerz, 2006).

## Conclusion

Here, we studied the feeding activity of the shallow-water sponge *H. panicea* and proposed a biosynthetic pathway (i.e., sponge elongases and desaturases) responsible for the biosynthesis of PLFAs. Furthermore, we assessed the potential of using deuterated water to measure the metabolic activity of suspension-feeding benthos. *H. panicea* had a condition index of 0.38 which indicates that this sponge species was likely starving in the Eastern Scheldt (Oosterschelde, Dutch Delta) where it might be in an inferior position compared to the large bivalve stocks. However, silicon uptake and oxygen consumption rates rather suggested that the sponges were well fed during the experiment, in fact, we provided them with a inactivated substrate bacteria that contained three orders of magnitude more C than they require to maintain their metabolism. Hence, our data were inconclusive.

Compound-specific stable isotope analyses of ^2^H and ^13^C in PLFA showed that the sponge microbiome uses exogenous C sources, in this case C from the inactivated substrate bacteria, to synthesize the bacteria-specific PLFAs *i*-C14:0, *ai*-C15:0, *i*-C17:0, and *ai*-C17:0. This *ai*-C15:0 is the precursor for enzymes of the sponge to build the long-chain PLFA *ai*-C25:2.

Other long-chain PLFAs (e.g., C24:1, C26:2, C26:3) are synthesized using C14:0 and C16:0 from exogenous sources.

Besides its application for the study of biosynthetic pathways, this pilot study proves that ^2^H from deuterated water can be used to measure the metabolic activity of marine sponges. However, we recommend to focus stronger on compound-specific ^2^H uptake rates than on bulk ^2^H uptake rates because of the considerable amount of ^2^H incorporation into inactive tissue. Additionally, we suggest to assess how ^2^H uptake rates differ between species with diverse biological traits or specimens maintained at different temperatures.

## Acknowledgements

Jan Peene, Pieter van Rijswijk, Peter van Breugel (all NIOZ-EDS), Jort Ossebar, and Ronald van Bommel (both NIOZ-MMB) are thanked for technical assistance during sample processing. Femke van Dam (Utrecht University) is thanked for assistance while conducting the experiments and field work and sponge taxonomist Rob van Soest is thanked who identified the sponge species. This study received funding by JPI Oceans – Impacts of deep-sea nodule mining project “Mining Impact 2” from the Dutch Research Council (NWO-ALW grant 856.18.003), from the research program NWO-Rubicon with project number 019.182EN.012, and from the NWO-Talent program Veni with project number VI.Veni.212.211.

## Supplementary Material

### Additional Information A1

Settings of the elemental analyzers (EAs) coupled to an isotope-ratio mass spectrometer (IRMS) to measure δ^13^C, δ^15^N, and δ^2^H in sponge tissue.

The EA gas settings for simultaneous measurements of **δ**^**13**^**C and δ**^**15**^**N** were 100 kPa He gas with 90 ml min^−1^ flow rate of carrier gas and 100 ml min^−1^ flow rate of reference gas, 280 kPa O_2_ gas with a 150 ml min^−1^ flow rate, and an injection end time of 3 sec. The operating parameters of the high-temperature reactor were a temperature of 1020°C in the left furnace and a temperature of 650°C in the right furnace. The oven with the gas chromatography column (HayeSep Q Packed GC Column, 80/100 mesh, length: 2 m, inner diameter: 2 mm; Supelco Inc, USA) had a temperature of 45°C. The interphase operated with a pressure of 0.7 kPa He, 0.7 kPa CO_2_ gas standard, and 0.7 kPa N_2_ gas standard.

For the measurements of δ^**2**^**H**, He gas served as carrier gas with 100 ml min^−1^ flow rate. The high-temperature reactor had a temperature of 1450°C in the left furnace and the oven with the gas chromatography column (O/H Separation column 1M 260 08240, length: 1 m; Elemental Microanalysis Ltd, UK) had a temperature of 50°C.

### Additional Information A2

Settings of the gas chromatograph (GC) with a flame ionization detector (FID) to measure C and H concentration of fatty acid methyl esters (FAMEs).

The GC-FID was equipped with a fused silica pre-column (length: 5 m, diameter: 0.32 mm) and a BPX70 column (length: 50 m length, inner diameter: 0.32 mm, film thickness: 0.25 μm; SGE Analytical Science). The injector had a base temperature of 50°C and injected the FAMEs in a splitless mode with a split flow of 25 ml min^−1^ and a splitless time of 2 min. The carrier gas He flowed with a constant flow of 2 min min^−1^.

| # | Rate ($^{\circ}\text{C min}^{-1}$) | Final temperature ($^{\circ}\text{C}$) | Hold time (min) |
| --- | --- | --- | --- |
| Initial |  | 50 | 3 |
| 1 | 30 | 110 | 5 |
| 2 | 2 | 220 | 0 |
| 3 | 20 | 240 | 5 |

### Additional Information A3

Settings of the GC coupled to a isotope ratio mass spectrometer (IRMS) to measure δ^13^C and δ^2^H in FAMES.

The GC-IRMS was equipped with a FS, deactivated pre-column (length: 1.5 m, diameter: 0.53 mm; Agilent, USA) and a BPX70 column (length: 50 m length, inner diameter: 0.32 mm, film thickness: 0.25 μm; SGE Analytical Science). For δ^**13**^**C** measurements, FAMEs were injected cold-on-column, and the carrier gas He flowed with a constant flow of 2 min min^−1^.

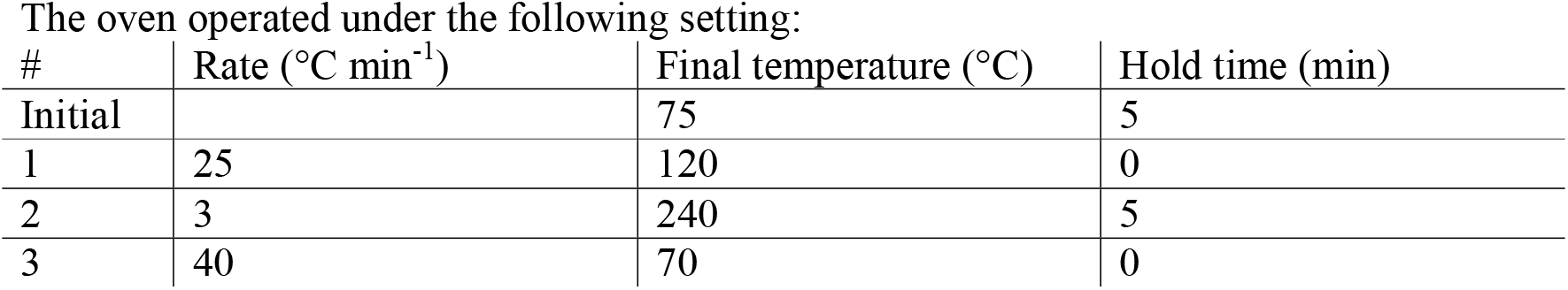

For δ^**2**^**H** measurements, FAMEs were injected cold-on-column, and the flow of the carrier gas (He) was 1.5 ml min^−1^.

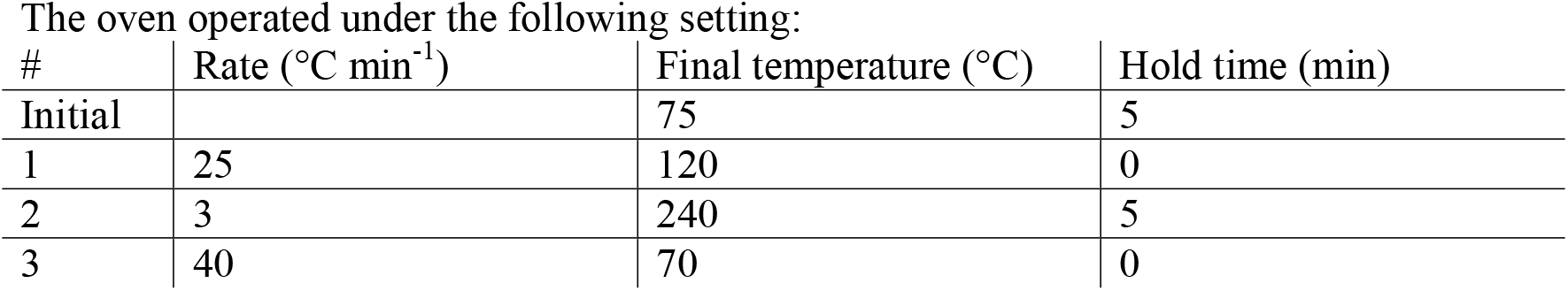

**Table S1.**
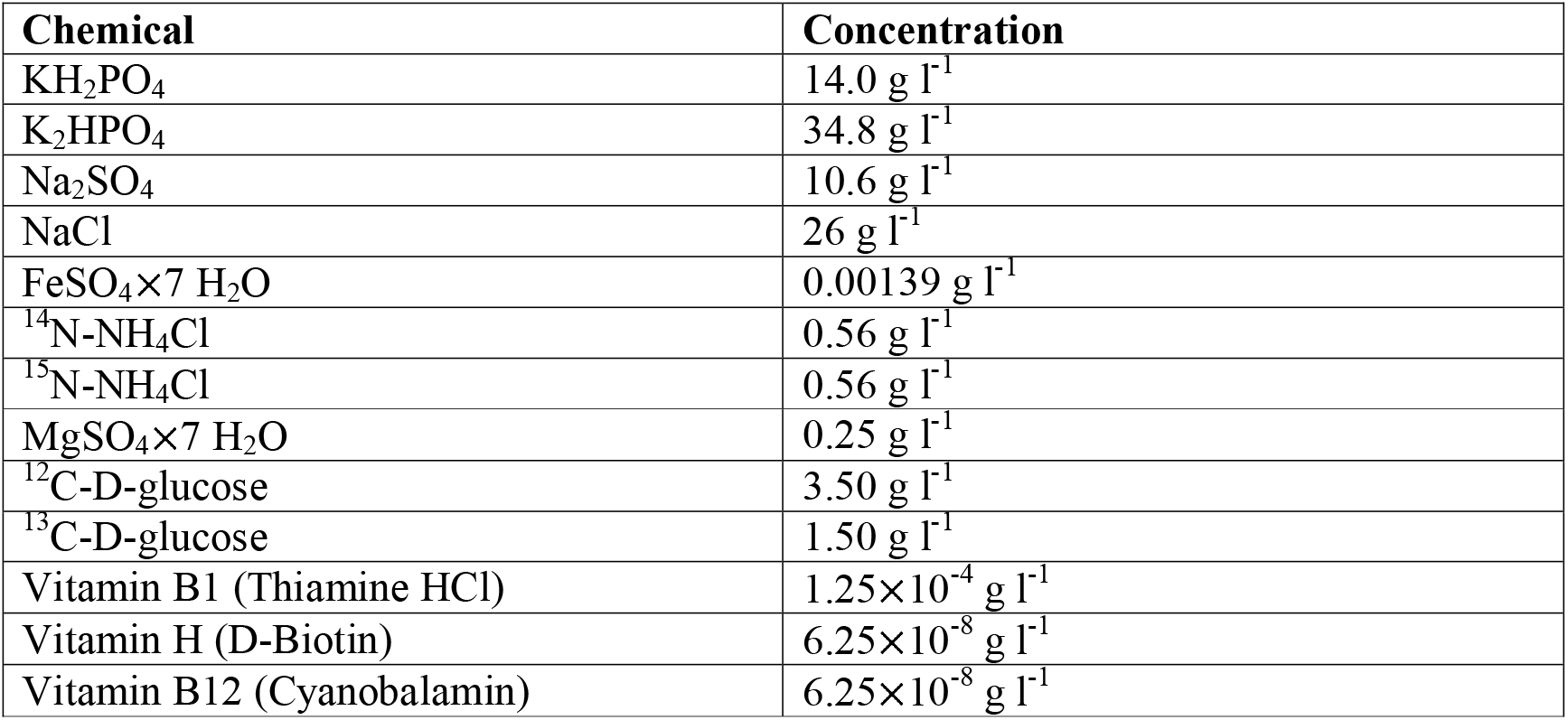
Culture medium for the substrate bacteria modified from the M63 minimum medium by Miller, (1972).

**Table S2.** Median, 5% – 95% range/ 25% – 75% range of δ^2^H_VSMOW-SLAP_-values of individual phospholipid-derived fatty acids (PLFAs) from *H. panicea* specimens with natural abundance stable isotopic composition (Nat. abundance), of PLFAs from dead sponge incubations (dead sponge), and from PLFAs of alive *H. panicea* (alive sponge). * 5^th^ – 95^th^ percentile ranges of δ^2^H values of PLFAs from nat. abundance sponges do not overlap with ranges of δ^2^H values of PLFAs from alive sponges. † 25^th^ – 75^th^ percentile ranges of δ^2^H values of PLFAs from nat. abundance sponges do not overlap with ranges of δ^2^H values of PLFAs from alive sponges. □ 5^th^ – 95^th^ percentile ranges of δ^2^H values of PLFAs from dead sponges do not overlap with ranges of δ^2^H values of PLFAs from alive sponges. ‡ 25^th^ – 75^th^ percentile ranges of δ^2^H values of PLFAs from dead sponges do not overlap with ranges of δ^2^H values of PLFAs from alive sponges.

| PLFA | Nat. abundance | Dead sponge | Alive sponge |
| --- | --- | --- | --- |
| i-C14:0*□†‡ | -318, -367 – -147/<br>-322 – -267 | -103, -285 – -88.1/<br>-285 – -88.1 | 221, 49.2 – 607/<br>113 – 432 |
| C14:0†‡ | -361, -422 – -318/<br>-405 – -338 | -374, -575 – -333/<br>-575 – -333 | -277, -368 – -225/<br>-316 – -247 |
| C14:1ω5c*□†‡ | -78.8, -166 – -58.7/<br>-91.7 – -68.5 | -51.7, -125 – -40.1/<br>-125 – -40.1 | 289, 79.2 – 523/<br>162 – 493 |
| ai-C15*□†‡ | -100, -330 – -43.4/<br>-276 – -70.1 | -76.7, -86.1 – -63.3/<br>-86.1 – -63.3 | 75.7, 8.77 – 166/<br>26.7 – 124 |
| C15:0□†‡ | -135, -158 – -70.3/<br>-149 – -114 | -149, -179 – -99.5/<br>-179 – -99.5 | -12.7, -94.6 – 147/<br>-44.4 – 81.3 |
| C15:1ω5c*□†‡ | -58.0, -92.3 – -49.9/<br>-71.6 – -56.1 | -57.9, -79.2 – -37.1/<br>-79.2 – -37.1 | 123, 51.4 – 306/<br>113 – 177 |
| C16:0*□†‡ | -119, -460 – -82.1/<br>-213 – -102 | -118, -297 – -86.2/<br>-297 – -86.2 | 183, 135 – 349/<br>159 – 207 |
| C16:1ω7c*□†‡ | -128, -148 – -111/<br>-129 – -116 | -110, -139 – -96.1/<br>-139 – -96.1 | 75.4, -7.42 – 226/<br>35.0 – 161 |
| i-C17:0*□†‡ | -116, -123 – -63.5/<br>-119 – -92.9 | -91.1, -120 – -70.2/<br>-120 – -70.2 | 104, -58.9 – 327/<br>34.2 – 177 |
| ai-C17:0*□†‡ | -71.5, -247 – -52.8/<br>-134 – -60.1 | -68.0, -90.4 – -59.5/<br>-90.4 – -59.5 | 96.3, 35.1 – 365/<br>84.1 – 202 |
| C17:0*†‡ | -140, -175 – -111/<br>-159 – -125 | -142, -202 – -95.2/<br>-202 – -95.2 | 13.0, -96.6 – 185/<br>-28.5 – 75.8 |
| C18:0*□†‡ | -183, -335 – -87.5/<br>-281 – -105 | -118, -152 – -98.2/<br>-152 – -98.2 | 79.3, -85.2 – 263/<br>48.5 – 101 |
| C18:1ω7c*□†‡ | -109, -181 – -93.1/<br>-129 – -95.3 | -158, -251 – -123/<br>-251 – -123 | 298, 21.3 – 588/<br>166 – 402 |
| C18:2ω(6,9t)*□†‡ | -145, -372 – -92.7/<br>-197 – -135 | -124, -191 – -96.9/<br>-191 – -96.9 | 865, 647 – 1105/<br>772 – 923 |
| C18:3ω(3,6,9) | -139, -169 – -71.2/<br>-156 – -101 | -99.2, -137 – -62.9/<br>-137 – -62.9 | 31.7, -209 – 478/<br>-106 – 105 |
| C18:3ω(6,9,12)†‡ | -191, -327 – -139/<br>-236 – -162 | -199, -273 – -155/<br>-273 – -155 | -59.6, -159 – 77.4/<br>-79.6 – -36.2 |
| C18:3ω4 | -95.3, -124 – -66.5/<br>-111 – -79.3 | -78.0, -130 – -41.4/<br>-130 – -41.4 | -40.0, -184 – 191/<br>-95.4 – 126 |
| C20:1 | -128, -189 – -88.9/<br>-150 – -117 | -151, -168 – -126/<br>-168 – -126 | -91.1, -192 – 150/<br>-150 – 33.0 |
| C20:1ω9c†‡ | -131, -175 – -101/<br>-134 – -126 | -130, -174 – -116/<br>-174 – -116 | -26.2, -161 – 184/<br>-89.6 – 18.8 |
| C20:4ω3†‡ | -61.7, -304 – -36.4/<br>-70.8 – -50.6 | -75.3, -114 – -50.8/<br>-114 – -50.8 | -27.3, -61.1 – 94.0/<br>-34.8 – -11.4 |
| C21:0□†‡ | -91.3, -280 – -73.3/<br>-101 – -76.6 | -109, -233 – -82.4/<br>-233 – -82.4 | 25.2, -64.1 – 321/<br>-9.17 – 131 |
| C22:6ω(3,6,9,12,15,18) | -121, -155 – -59.9/<br>-133 – -109 | -63.0, -120 – -28.7/<br>-120 – -28.7 | -69.7, -183 – 379/<br>-118 – 241 |
| C23:0†‡ | -157, -370 – -130/<br>-191 – -137 | -182, -219 – -149/<br>-219 – -149 | -98.1, -165 – 398/<br>-101 – -54.9 |
| C24:0† | -87.3, -125 – -34.4/<br>-117 – -86.8 | -76.5, -158 – -52.2/<br>-158 – -52.2 | -40.3, -105 – 284/<br>-55.3 – 107 |
| C24:1†‡ | -114, -137 – -102/<br>-123 – -109 | -138, -148 – -97.4/<br>-148 – -97.4 | 293, -107 – 1365/<br>200 – 1063 |
| (i-/ ai-C25:2)*□†‡ | -119, -191 – -80.2/<br>-136 – -95.6 | -113, -145 – -87.9/<br>-145 – -87.9 | 128, -7.16 – 351/<br>50.0 – 269 |
| i-C25:2*□†‡ | -177, -225 – -130/<br>-212 – -142 | -155, -176 – -112/<br>-176 – -112 | -7.28, -70.8 – 150/<br>-39.9 – 29.0 |
| ai-C25:2†‡ | -205, -263 – -182/<br>-237 – -186 | -209, -259 – -157/<br>-259 – -157 | -103, -222 – 88.1/<br>-132 – -61.3 |
| C25:2*□†‡ | -120, -175 – -92.0/<br>-126 – -112 | -130, -154 – -105/<br>-154 – -105 | -8.56, -63.7 – 335/<br>-55.6 – 126 |
| C26:2*□†‡ | -142, -150 – -135/<br>-146 – -140 | -143, -148 – -118/-<br>148 – -118 | -39.4, -114 – 358/<br>-70.6 – 57.0 |
| C26:3*□†‡ | -161, -208 – -134/<br>-163 – -156 | -163, -165 – -141/<br>-165 – -141 | -33.2, -109 – 559/<br>-97.9 – 219 |
| C27:2†‡ | -140, -232 – -130/<br>-183 – -136 | -145, -188 – -141/<br>-188 – -141 | -22.5, -174 – 1867/<br>-116 – 166 |

**Figure S1.**
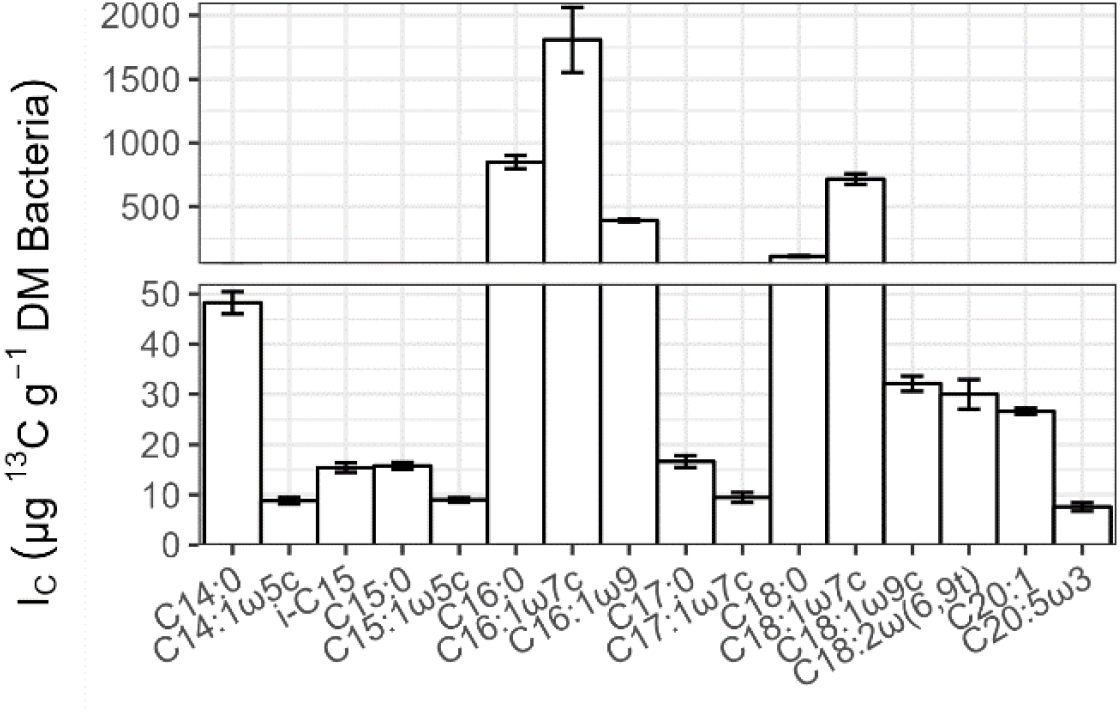
Incorporation of ^13^C (I_C_) into phospholipid-derived fatty acids (PLFA) (μg ^13^C g^−1^ dry mass bacteria) from the substrate bacteria that was fed to the alive sponges. Error bars represent 1 SE.

**Figure S2.**
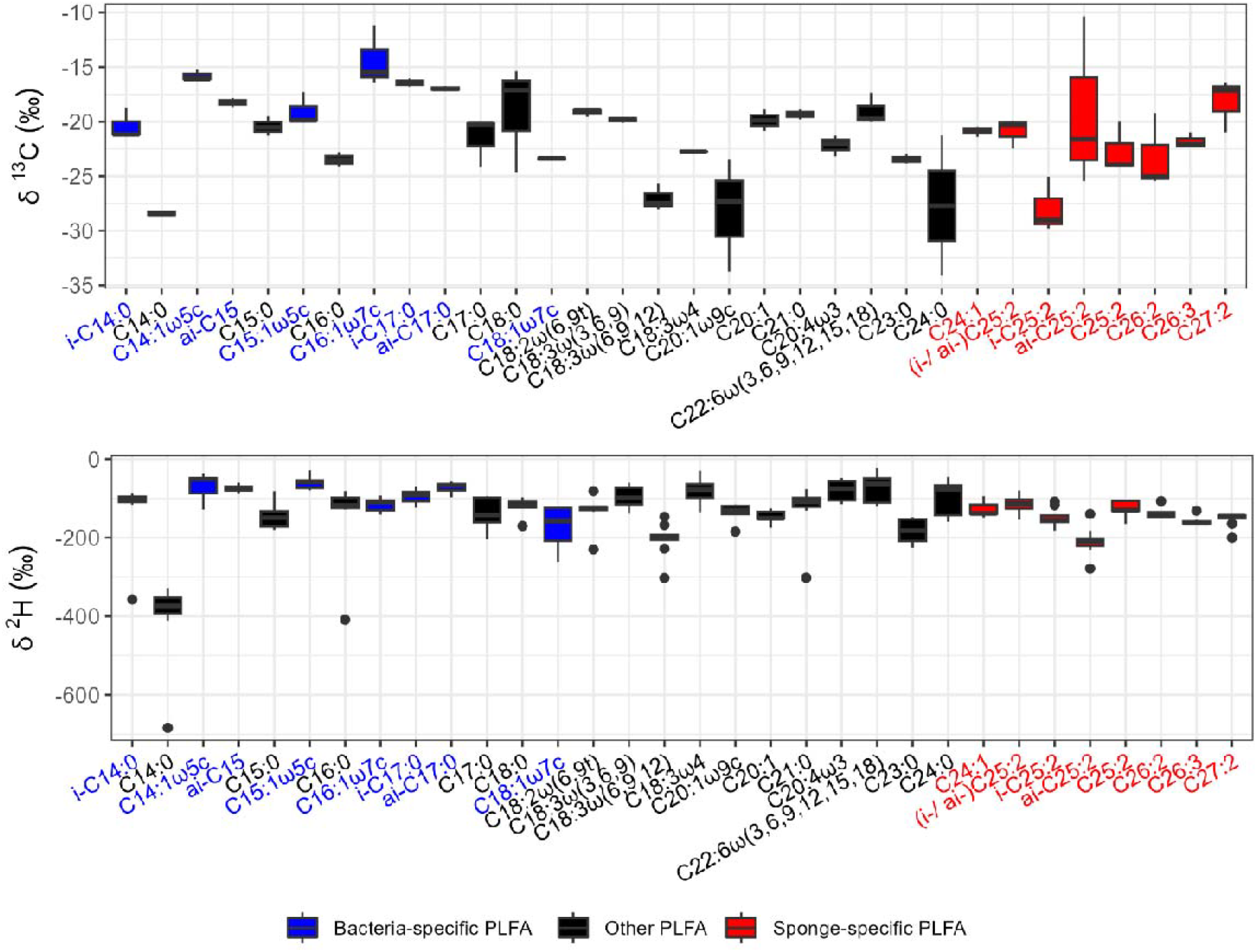
Stable isotopic enrichment of PLFAs extracted from *H. panicea* specimens from dead-sponge incubations. (Upper panel) δ^13^C_VPDB_ values (‰) of PLFAs and (lower panel) δ^2^H_VSMOW-SLAP_ values (‰) of PLFAs.

**Figure S3.**
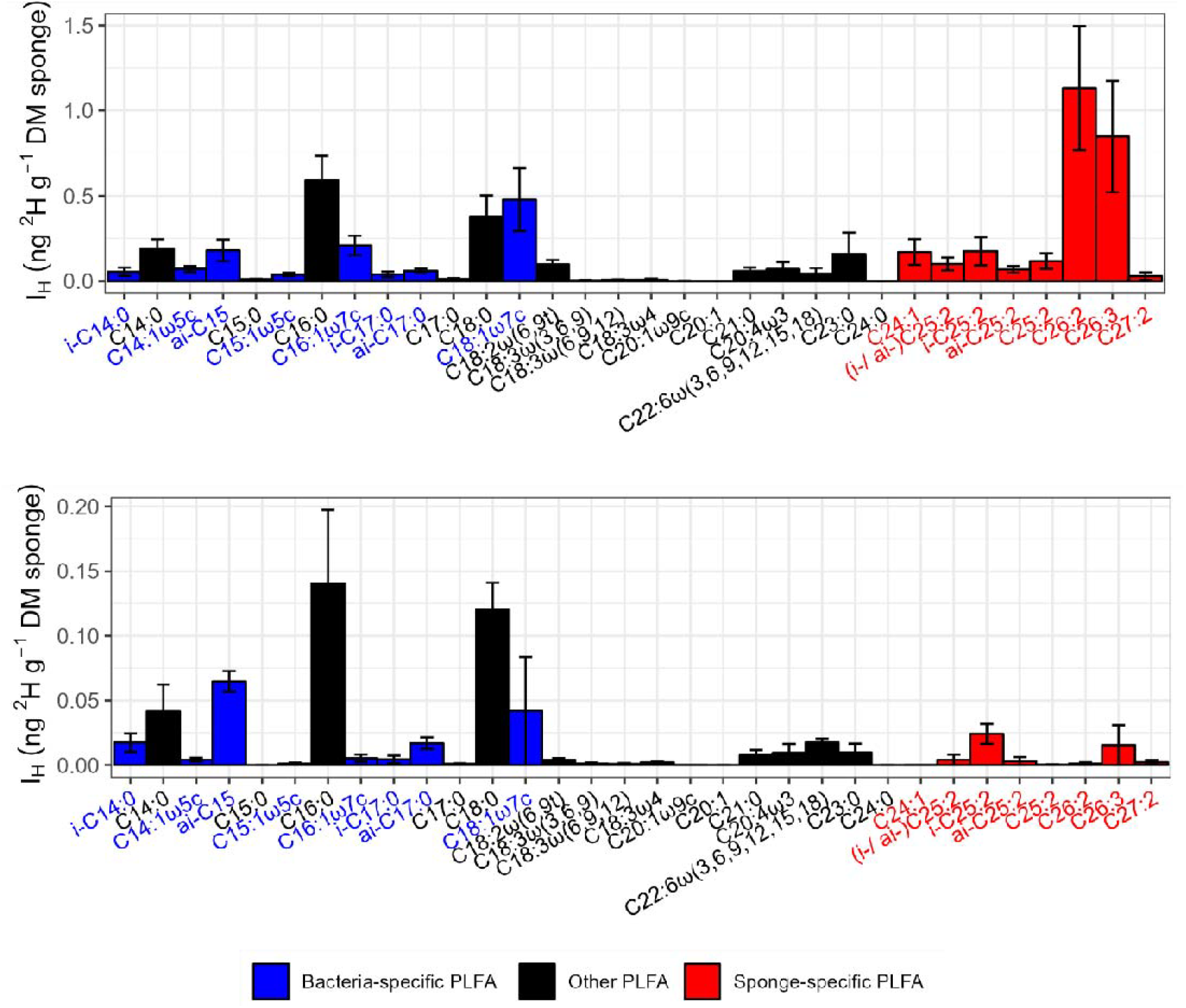
Incorporation of ^2^H into phospholipid-derived fatty acids (PLFA) (ng ^2^H g^−1^ dry mass sponge) from incubations of alive sponges (upper panel) and incubations of dead sponges (lower panel). Error bars indicate 1 SE.

